# S/HIC: Robust identification of soft and hard sweeps using machine learning

**DOI:** 10.1101/024547

**Authors:** Daniel R. Schrider, Andrew D. Kern

## Abstract

Detecting the targets of adaptive natural selection from whole genome sequencing data is a central problem for population genetics. However, to date most methods have shown sub-optimal performance under realistic demographic scenarios. Moreover, over the past decade there has been a renewed interest in determining the importance of selection from standing variation in adaptation of natural populations, yet very few methods for inferring this model of adaptation at the genome scale have been introduced. Here we introduce a new method, S/HIC, which uses supervised machine learning to precisely infer the location of both hard and soft selective sweeps. We show that S/HIC has unrivaled accuracy for detecting sweeps under demographic histories that are relevant to human populations, and distinguishing sweeps from linked as well as neutrally evolving regions. Moreover we show that S/HIC is uniquely robust among its competitors to model misspecification. Thus even if the true demographic model of a population differs catastrophically from that specified by the user, S/HIC still retains impressive discriminatory power. Finally we apply S/HIC to the case of resequencing data from human chromosome 18 in a European population sample and demonstrate that we can reliably recover selective sweeps that have been identified earlier using less specific and sensitive methods.

## INTRODUCTION

The availability of population genomic data has empowered efforts to uncover the selective, demographic, and stochastic forces driving patterns of genetic variation within species. Chief among these are attempts to uncover the genetic basis of recent adaptation [1]. Indeed, recent advances in genotyping and sequencing technologies have been accompanied by a proliferation of statistical methods for identifying recent positive selection [see 2 for recent review].

Most methods for identifying positive selection search for the population genetic signature of a “selective sweep” [3], wherein the rapid fixation of a new beneficial allele leaves a valley of diversity around the selected site [4–6], about which every individual in the population exhibits the same haplotype (i.e. the genetic background on which the beneficial mutation occurred). At greater genetic distances, polymorphism recovers as recombination frees linked neutral variants from the homogenizing force of the sweep [4]. This process also produces an excess of low- and high-frequency derived alleles [7, 8], and increased allelic association, or linkage disequilibrium (LD), on either side of the sweep [9], but not across the two flanks of the sweep [10, 11]. Selective fixation of *de novo* beneficial mutations such as described by Maynard Smith and Haigh [5] are often referred to as “hard sweeps.”

More recently, population geneticists have begun to consider the impact of positive selection on previously standing genetic variants [12, 13]. Under this model of adaptation, an allele initially evolves under drift for some time, until a change in the selective environment causes it to confer a fitness advantage and sweep to fixation. In contrast to the hard sweep model, the selected allele is present in multiple copies prior to the sweep. Thus, because of mutation and recombination events occurring near the selected site during the drift phase, the region containing this site may exhibit multiple haplotypes upon fixation [14]. The resulting reduction in diversity is therefore less pronounced than under the hard sweep model [12, 15]. For this reason sweeps from standing genetic variation are often referred to as “soft sweeps.” Soft sweeps will not skew the allele frequencies of linked neutral polymorphisms toward low and high frequencies to the same extent as hard sweeps [16], and may even present an excess of intermediate frequencies [17]. This mode of selection will also have a different impact on linkage disequilibrium: LD will be highest at the target of selection rather than in flanking regions [18]. In very large populations, selection on mutations that are immediately beneficial may also produce patterns of soft sweeps rather than hard sweeps, as the adaptive allele may be introduced multiple times via recurrent mutation before the sweep completes [14, 19]. While this model of a soft sweep is similar to that of selection on standing variation in that it will produce additional haplotypes carrying the selected allele, there are differences in the patterns of polymorphism produced by these two types of soft sweeps [18, 20]. Here, we examine only the model of selection on a single standing variant.

Adaptation could proceed primarily through selection on standing variation if the selective environment shifts frequently relative to the time scale of molecular evolution, and if there is enough standing variation segregating in the population on which selection may act following such a shift [12, 21]. However, it is important to note that selection on standing variation may produce a hard sweep of only one haplotype containing the adaptive mutation if this allele is present at low enough frequency prior to sweep [16, 22]. In other words, the observation of hard sweeps may be consistent with selection on standing variation as well as selection on *de novo* mutations. For these and other reasons, there is some controversy over whether adaptation will result in soft sweeps in nature [22]. This could be resolved by methods that can accurately discriminate between hard and soft sweeps. To this end, some recently devised methods for detecting population genetic signatures of positive selection consider both types of sweeps [23–25]. Unfortunately, it may often be difficult to distinguish soft sweeps from regions flanking hard sweeps due to the “soft shoulder” effect [18].

Here we present a method that is able to accurately distinguish between hard sweeps, soft sweeps on a single standing variant, regions linked to sweeps (or the “shoulders” of sweeps), and regions evolving neutrally. This method incorporates spatial patterns of a variety of population genetic summary statistics across a large genomic window in order to infer the mode of evolution governing a focal region at the center of this window. We combine many statistics used to test for selection using an Extremely Randomized Trees classifier [26], a powerful supervised machine learning classification technique. We refer to this method as Soft/Hard Inference through Classification (S/HIC, pronounced “shick”). By incorporating multiple signals in this manner S/HIC achieves inferential power exceeding that of any individual test. Furthermore, by using spatial patterns of these statistics within a broad genomic region, S/HIC is able to distinguish selective sweeps not only from neutrality, but also from linked selection with much greater accuracy than other methods. Thus, S/HIC has the potential to identify more precise candidate regions around recent selective sweeps, thereby narrowing down searches for the target locus of selection. Further, S/HIC’s reliance on large-scale spatial patterns makes it more robust to non-equilibrium demography than previous methods, even if the demographic model is misspecified during training. This is vitally important, as the true demographic history of a population sample may be unknown. Finally, we demonstrate the utility of our approach by applying it to chromosome 18 in the CEU sample from the 1000 Genomes dataset [27], recovering most of the sweeps identified previously in this population through other methods; we also highlight a compelling novel candidate sweep in this population.

## METHODS

### Supervised machine learning to detect soft and hard sweeps

We sought to devise a method that could not only accurately distinguish among hard sweeps, soft sweeps, and neutral evolution, but also among these modes of evolution and regions linked to hard and soft sweeps, respectively [18]. Such a method would not only be robust to the soft shoulder effect, but would also be able to more precisely delineate the region containing the target of selection by correctly classifying unselected but closely linked regions. In order to accomplish this, we sought to exploit the impact of positive selection on spatial patterns of several aspects of variation surrounding a sweep. Not only will a hard sweep create a valley of diversity centered around a sweep, but it will also create a skew toward high frequency derived alleles flanking the sweep and intermediate frequencies at further distances [7, 8], reduced haplotypic diversity at the sweep site [24], and increased LD along the two flanks of the sweep but not between them [10]. For soft sweeps, these expected patterns may differ considerably [14, 16, 18], but also depart from the neutral expectation.

While some of these patterns of variation have been used individually for sweep detection [e.g. 10, 28], we reasoned that by combining spatial patterns of multiple facets of variation we would be able to do so more accurately. To this end, we designed a machine learning classifier that leverages spatial patterns of a variety of population genetic summary statistics in order to infer whether a large genomic window recently experienced a selective sweep at its center. We accomplished this by partitioning this large window into adjacent subwindows, measuring the values of each summary statistic in each subwindow, and normalizing by dividing the value for a given subwindow by the sum of values for this statistic across all subwindows within the same window to be classified. Thus, for a given summary statistic, *x* we used the following vector:

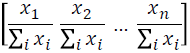

where the larger window has been divided into *n* subwindows, and *x_i_* is the value of the summary statistic *x* in the *i^th^* subwindow. Thus, this vector captures differences in the relative values of a statistic across space within a large genomic window, but does not include the actual values of the statistic. In other words, this vector captures only the shape of the curve of the statistic *x* across the large window that we wish to classify. Our goal is to then infer a genomic region’s mode of evolution based on whether the shapes of the curves of various statistics surrounding this region more closely resemble those observed around hard sweeps, soft sweeps, neutral regions, or loci linked to hard or soft sweeps. In addition to allowing for discrimination between sweeps and linked regions, this strategy was motivated by the need for accurate sweep detection in the face of a potentially unknown nonequilibrium demographic history, which may grossly affect values of these statistics but may skew their expected spatial patterns to a much lesser extent. In total, we constructed these vectors for each of π [29], 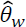[30], 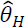[8] the number of distinct haplotypes, average haplotype homozygosity, *H*_12_ and *H*_2_/*H*_1_ [24, 31], *Z_nS_* [9], and the maximum value of *ω* [10]. Thus, we represent each large genomic window by the following vector, to which we refer as the feature vector:

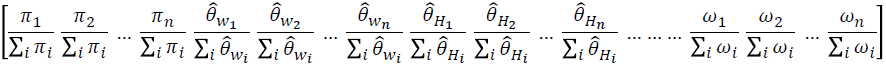

We sought to discriminate between hard sweeps, regions linked to hard sweeps, soft sweeps, regions linked to soft sweeps, and neutrally evolving regions on the basis of the values of the vectors defined above. While Berg and Coop [20] recently derived approximations for the site frequency spectrum (SFS) for a soft sweep under equilibrium population size, and *π*, the joint probability distribution of the values all of the above statistics at varying distances from a sweep is unknown. Moreover expectations for the SFS surrounding sweeps (both hard and soft) under nonequilibrium demography remain analytically intractable. Thus rather than taking a likelihood approach, we opted to use a supervised machine learning framework, wherein a classifier is trained from simulations of regions known to belong to one of these five classes. We trained an Extra-Trees classifier (aka extremely randomized forest; [26]) from coalescent simulations (described below) in order to classify large genomic windows as experiencing a hard sweep in the central subwindow, a soft sweep in the central subwindow, being closely linked to a hard sweep, being closely linked to a soft sweep, or evolving neutrally according to the values of its feature vector (Fig 1).

**Fig. 1.**
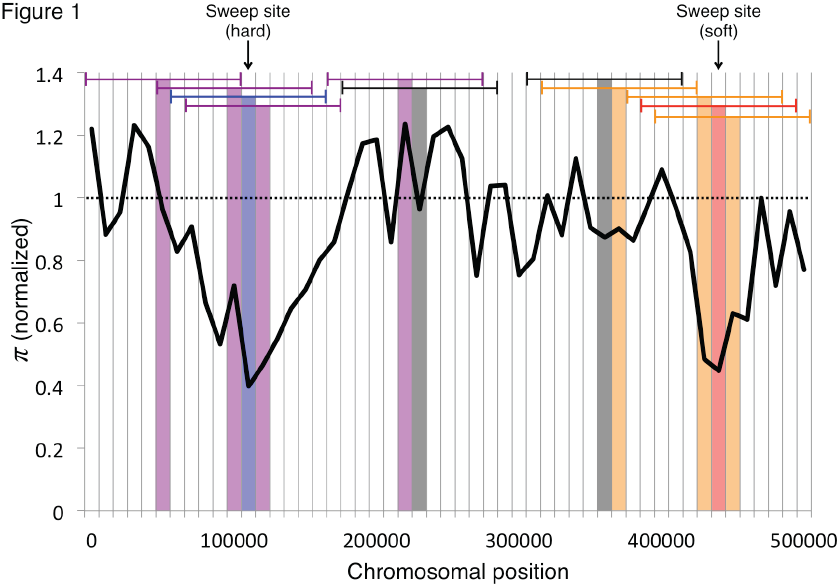
Examples of the five classes used by S/HIC. S/HIC classifies each window as a hard sweep (blue), linked to a hard sweep (purple), a soft sweep (red), linked to a soft sweep (orange), or neutral (gray). This classifier accomplishes this by examining values of various summary statistics in 11 different windows in order to infer the mode of evolution in the central window (the horizontal blue, purple, red, orange, and gray brackets). Regions that are centered on a hard (soft) selective sweep are defined as hard (soft). Regions that are not centered on selective sweeps but have their diversity impacted by a hard (soft) selective sweep but are not centered on the sweep are defined as hard-linked (soft-linked). Remaining windows are defined as neutral. S/HIC is trained on simulated examples of these five classes in order to distinguish selective sweeps from linked and neutral regions in population genomic data.

Briefly, the Extra-Trees classifier is an ensemble classification technique that harnesses a large number classifiers referred to as decision trees. A decision tree is a simple classification tool that uses the values of multiple features for a given data instance, and creates a branching tree structure where each node in the tree is assigned a threshold value for a given feature. If a given data point’s (or instance’s) value of the feature at this node is below the threshold, this instance takes the left branch, and otherwise it takes the right. At the next lowest level of the tree, the value of another feature is examined. When the data instance reaches the bottom of the tree, it is assigned a class inference based on which leaf it has landed [32]. Typically, a decision tree is built according to an algorithm designed to optimize its accuracy [32]. The Extra-Trees classifier, on the other hand, builds a specified number of semi-randomly generated decision trees. Classification is then performed by simply taking the class receiving the most “votes” from these trees [26], building on the strategy of random forests [33]. While individual decision trees may be highly inaccurate, the practice of aggregating predictions from many semi-randomly generated decision trees has been proved to be quite powerful [34].

In the following sections we describe our methodology for training, testing, and applying our Extra-Trees classifier for identifying positive selection.

### Coalescent simulations for training and testing

We simulated data for training and testing of our classifier using our coalescent simulator, discoal_multipop (https://github.com/kern-lab/discoal_multipop). As discussed in the Results, we simulated training sets with different demographic histories (Table S1), and, for positively selected training examples, different ranges of selection coefficients (*α*=2*Ns*, where *s* is the selective advantage and *N* is the population size). For each combination of demographic history and range of selection coefficients, we simulated large chromosomal windows that we later subdivided into 11 adjacent and equally sized subwindows. We then simulated training examples with a hard selective sweep whose selection coefficient was uniformly drawn from the specified range, *U*(*α*_low_, *α*_high_). We generated 11,000 sweeps: 1000 where the sweep occurred in the center of the leftmost of the 11 subwindows, 1000 where the sweep occurred in the second subwindow, and so on. We repeated this same process for soft sweeps at each location; these simulations had an additional parameter, the derived allele frequency, *f*, at which the mutation switches from evolving under drift to sweeping to fixation, which we drew from *U*(0.05, 0.2), *U*(2/2*N*, 0.05), or *U*(2/2*N*, 0.2) as described in the Results. For our equilibrium demography scenario, we drew the fixation time of the selective sweep from *U*(0, 0.2)×*N* generations ago, while for non-equilibrium demography the sweeps completed more recently (see below). We also simulated 1000 neutrally evolving regions. Unless otherwise noted, for each simulation the sample size was set to 100 chromosomes.

For each combination of demographic scenario and selection coefficient, we combined our simulated data into 5 equally-sized training sets (Fig 1): a set of 1000 hard sweeps where the sweep occurs in the middle of the central subwindow (i.e. all simulated hard sweeps); a set of 1000 soft sweeps (all simulated soft sweeps); a set of 1000 windows where the central subwindow is linked to a hard sweep that occurred in one of the other 10 windows (i.e. 1000 simulations drawn randomly from the set of 10000 simulations with a hard sweep occurring in a non-central window); a set of 1000 windows where the central subwindow is linked to a soft sweep (1000 simulations drawn from the set of 10000 simulations with a flanking soft sweep); and a set of 1000 neutrally evolving windows unlinked to a sweep. We then generated a replicate set of these simulations for use as an independent test set.

### Training the Extra-Trees classifier

We used the python scikit-learn package (http://scikit-learn.org/) to train our Extra-Trees classifier and to perform classifications. Given a training set, we trained our classifier by performing a grid search of multiple values of each of the following parameters: max_features (the maximum number of features that could be considered at each branching step of building the decision trees, which was set to 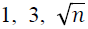, or *n*, where *n* is the total number of features); max_depth (the maximum depth a decision tree can reach; set to 3, 10, or no limit), min_samples_split (the minimum number of training instances that must follow each branch when adding a new split to the tree in order for the split to be retained; set to 1, 3, or 10); min_samples_leaf. (the minimum number of training instances that must be present at each leaf in the decision tree in order for the split to be retained; set to 1, 3, or 10); bootstrap (a binary parameter that governs whether or not a different bootstrap sample of training instances is selected prior to the creation of each decision tree in the classifier); criterion (the criterion used to assess the quality of a proposed split in the tree, which is set to either Gini impurity [35] or to information gain, i.e. the change in entropy [32]). The number of decision trees included in the forest was always set to 100. After performing a grid-search with 10-fold cross validation in order to identify the optimal combination of these parameters, we used this set of parameters to train the final classifier.

We used the scikit-learn package to assess the importance of each feature in our Extra-Trees classifiers. This is done by measuring the mean decrease in Gini impurity, multiplied by the average fraction of training samples that reach that feature across all decision trees in the classifier. The mean decrease in impurity for each feature is then divided by the sum across all features to give a relative importance score, which we show in Table S2. We also show values of Extra-Trees classifier parameters resulting from grid searchers in Table S3.

### Comparisons with other methods

We compared the performance of our classifier to that of various other methods. First, we examined two population genetic summary statistics: Tajima’s *D* [36] and Kim and Nielsen’s *ω_max_* [10] (which we refer to as *ω* for simplicity), calculating their values in each subwindow within each large simulated chromosome that we generated for testing (see above). We also used Nielsen et al.’s composite-likelihood ratio test, referred to as CLR or SweepFinder [28], which searches for the spatial skew in allele frequencies expected surrounding a hard selective sweep. When testing SweepFinder’s ability to discriminate between modes of evolution within larger regions, we computed the composite-likelihood ratio between the sweep and neutral models at 200 sites across each of the 11 subwindows of our large simulated test regions, taking the maximum CLR value. The only training necessary for SweepFinder was to specify the neutral site frequency spectrum.

Next, we used scikit-learn to implement Ronen et al.’s [37] *SFselect*, a support vector machine classifier that discriminates between selection and neutrality on the basis of a region’s binned and weighted SFS. In our implementation we collapsed the SFS into 10 bins as suggested by Ronen et al., and also added soft sweeps as a third class (in addition to hard sweeps and neutrality), using Knerr et al.’s [38] method for extending a binary classifier to perform multi-class classification. We trained this classifier from simulated data following the same demographic and selective scenarios used to train our own classifier, and with the same number of simulated training instances, but these simulations encapsulated much smaller regions (equivalent to the size of one of our eleven subwindows). To avoid confusion with the original *SFselect*, which only handles hard sweeps, we refer to this implementation as *SFselect+*. For further comparisons, we also trained a support vector machine using a vector of two statistics: the maximum values of the SweepFinder CLR statistic and *ω* (a subset of the features in the Pavlidis et al.’s SVM [39]). We refer to this method as CLR+*ω*, and trained it in the same manner as *SFselect+*, except for the different feature vector.

We also tested the performance of the evolBoosting [40], an R package which uses an machine learning approach called boosting [41] to classify genomic windows as sweeps or neutral on the basis of several statistics, including Tajima’s *D*, Fay and Wu’s *H* [8], integrated haplotype homozygosity (*iHH*; [42]), and several others. Boosting was also recently used by Pybus et al. [43] to accurately detect hard and partial sweeps and make coarse inferences about sweep ages. Like S/HIC, this method uses a vector of the values of each of these statistics from several subwindows surrounding the region being classified. However, unlike S/HIC, this method does not take the relative values of these statistics in each subwindow divided by the sum across all subwindows, instead just taking the value of the statistic measured in that subwindow. As with *SFselect*, we extend this method to discriminate between hard sweeps, soft sweeps, and neutral windows. This was done by first training a classifier to distinguish between sweeps (hard and soft, balanced in number within the training set) from neutral windows and secondarily training a classifier to distinguish between hard and soft sweeps.

Finally, we implemented a version of Garud et al.’s [24] scan for hard and soft sweeps. Garud et al.’s method uses an Approximate Bayesian Computation-like approach to calculate Bayes Factors to determine whether a given region is more similar to a hard sweep or a soft sweep by performing coalescent simulations. For this we performed simulations with the same parameters as we used to train *SFselect+*, but generated 100,000 simulations of each scenario in order to ensure that there was enough data for rejection sampling. We then used two statistics to summarize haplotypic diversity within these simulated data: *H*_12_ and *H*_2_/*H*_1_ [31]. All simulated regions whose vector [*H*_12_ *H*_2_*/H*_1_] lies within a Euclidean distance of 0.1 away from the vector corresponding to the data instance to be classified are then counted [24]. The ratio of simulated hard sweeps to simulated soft sweeps within this distance cutoff is then taken as the Bayes Factor. Note that Garud et al. restricted their analysis of the *D. melanogaster* genome to only the strongest signals of positive selection, asking whether they more closely resembled hard or soft sweeps. Therefore when testing the ability of Garud et al.’s method to distinguish selective sweeps from both linked and neutrally evolving regions, we used large simulated windows and simply examined the value of *H*_12_ within the subwindow that exhibited the largest value in an effort to mimic their strategy of using *H*_12_ peaks [24].

We summarized each method’s power using the receiver operating characteristic (ROC) curve, making these comparisons for the following binary classification problems: discriminating between hard sweeps and neutrality, between hard sweeps and soft sweeps, between selective sweeps (hard or soft) and neutrality, and between selective sweeps (hard or soft) and unselected regions (including both neutrally evolving regions and regions linked to selective sweeps). For each of these comparisons we constructed a balanced test set with a total of 1000 simulated regions in each class, so that the expected accuracy of a completely random classifier was 50%, and the expected area under the ROC curve (AUC) was 0.5. Whenever the task involved a class that was a composite of two or more modes of evolution, we ensured that the test set was comprised of equal parts of each subclass. For example, in the selected (hard or soft) versus unselected (neutral or linked selection) test, the selected class consisted of 500 hard sweeps and 500 soft sweeps, while the unselected class consisted of 333 neutrally evolving regions, 333 regions linked to hard sweeps, and 333 regions linked to soft sweeps (and one additional simulated region from one of these test sets randomly selected, so that the total size of the unselected test set was 1000 instances). As with our training sets, we considered the true class of a simulated test region containing a hard (soft) sweep occurring in any but the central subwindow to be hard-linked (soft-linked)—even if the sweep occurred only one subwindow away from the center.

The ROC curve is generated by measuring performance at increasingly lenient thresholds for discriminating between the two classes. We therefore required each method to output a real-valued measure proportional to its confidence that a particular data instance belongs the first of the two classes. For S/HIC, we used the posterior classification probability from the Extra-Trees classifier obtained using scikit-learn’s predict_proba method. For *SFselect+*, we used the value of the SVM decision function. For SweepFinder, we used the composite likelihood ratio. For Garud et al.’s method, we used the fraction of accepted simulations (i.e. within a Euclidean distance of 0.1 from the test instance) that were of the first class: for example, for hard vs. soft, this is the number of accepted simulations that were hard sweeps divided by the number of accepted simulations that were either hard sweeps or soft sweeps. For Tajima’s *D* [36] and Kim and Nielsen’s *ω* [10], we simply used the values of these statistics.

### Simulating sweeps under non-equilibrium demographic models

To examine the power and sensitivity of S/HIC under non-equilibrium demographic histories, we simulated training and test datasets from a few scenarios that might be relevant to researchers. Firstly we examined the power of our method under two complex population size histories that are relevant to humans. Secondly we examined the case of simple population bottlenecks, as might be common in populations that have recently colonized new locales, using two levels of bottleneck severity.

We simulated training and test datasets from Tennessen et al.’s [44] European demographic model (Table S1). This model parameterizes a population contraction associated with migration out of Africa, a second contraction followed by exponential population growth, and a more recent phase of even faster exponential growth. Values of *θ* and *ρ*=4*Nr* were drawn from prior distributions (Table S1), allowing for variation within the training data, whose means were selected from recent estimates of human mutation [45] and recombination rates [46], respectively. For simulations with selection, we drew values of *α* from 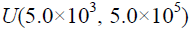, and drew the fixation time of the sweeping allele form *U*(0, 51,000) years ago (i.e. the sweep completed after the migration out of Africa).

We also generated simulations of Tennessen et al.’s African demographic model, which consists of exponential population growth beginning ∼5,100 years ago (Table S1). We generated two sets of these simulations: one where *α* was drawn from 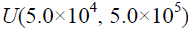, and one with *α* drawn from 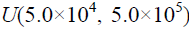. The sample size of these simulated data sets was set to 100 chromosomes. These two sets were then combined into a single training set. For these simulations, the sweep was constrained to complete some time during the exponential growth phase (no later than 5,100 years ago).

Finally, we examined two models with a population size bottleneck. The first was taken from Thornton and Andolfatto [47], and models the demographic history of a European population sample of *D. melanogaster* (Table S1). This model consists of a population size reduction 0.044×2*N* generations ago to 2.9% of the ancestral population size, and then 0.0084×2*N* generations ago the population recovers to its original size. The second bottleneck model we used was identical except the population contraction was less severe (reduction to 29% of the ancestral population size). For sweep simulations under both of these bottleneck scenarios, we drew *α* from 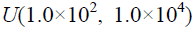. For all of our non-equilibrium demographic histories, when simulating soft sweeps on a previously standing variant, we drew the derived allele frequency at the onset of positive selection from *U*(2/2*N*, 0.2). For each demographic model in Table S1, we show in Fig S1 the means and standard deviations of Tajima’s *D* across 11 windows at increasing distances from a selective sweep (for one possible sweep scenario), as well as values from neutrally evolving windows for comparison.

For each demographic model in Table S1, we show in Fig S1 the means and standard deviations of Tajima’s *D* across 11 windows at increasing distances from a selective sweep (for one possible sweep scenario), as well as values from neutrally evolving windows for comparison.

### Application to human chromosome 18 from the 1000 Genomes CEU sample

We applied our method to chromosome 18 from the Phase I data release from the 1000 Genomes project [27]. We restricted this analysis to the CEU population sample (individuals with European ancestry, sampled from Utah), and trained S/HIC using data from the European demographic model described above. After training this classifier, we prepared data from chromosome 18 in CEU for classification. Prior to constructing feature vectors, we first performed extensive filtering for data quality. First, we masked all sites flagged by the 1000 Genomes Project as being unfit for population genetic analyses due to having either limited or excessive read-depth or poor mapping quality (according to the strictMask files for the Phase I data set which are available at http://www.1000genomes.org/). In order to remove additional sites lying within repetitive sequence wherein genotyping may be hindered, we eliminated sites with 50 bp read mappability scores less than one [48] and also sites masked by RepeatMasker (http://www.repeatmasker.org). Finally, we attempted to infer the ancestral state at each remaining site, using the chimpanzee [49] and macaque [50] genomes as outgroups. For each site, if the chimpanzee and macaque genomes agreed, we used this nucleotide as our inferred ancestral state. If instead only the chimpanzee or the macaque genome had a nucleotide aligned to the site, we used this base as our inferred ancestral state. For sites that were SNPs, we also required that the inferred ancestral state matched one of the two human alleles. For all cases where these criteria were not met, we discarded the site.

After data filtering, we calculated summary statistics within adjacent 200 kb windows across the entire chromosome. Importantly, we divided the values of each summary statistic by the number of sites in the window, ignoring sites filtered as discussed above (i.e. *π* summarizes average nucleotide diversity per site rather than total diversity in the subwindow). Windows with >50% of sites removed during the filtering processes were omitted from our analysis. These two steps limited the effect of variation in the number of unfiltered sites from window to window our classification. For the remaining windows, we used a sliding window approach with a 2.2 Mb window and a 200 kb step size to calculate the feature vector in the same manner as for our simulated data, and then applied S/HIC to this feature vector to infer whether the central subwindow of this 2.2 Mb region contained a hard sweep, a soft sweep, was linked to a hard sweep, linked to a soft sweep, or evolving neutrally. Visualization of candidate regions was performed using the UCSC Genome Browser [51]. We used hg19 coordinates for all of our analyses using human data.

### Software availability

Our classification tool is available at https://github.com/kern-lab/shIC, along with software for generating the feature vectors used in this paper (either from simulated training data or from real data for classification).

## RESULTS

### S/HIC accurately detects hard sweeps

The most basic task that a selection scan must be able to perform is to distinguish between hard sweeps and neutrally evolving regions, as the expected patterns of nucleotide diversity, haplotypic diversity, and linkage disequilibrium produced by these two modes of evolution differ dramatically [5, 8, 10, 18, 24, 52]. We therefore begin by comparing S/HIC’s power to discriminate between hard sweeps and neutrality to that of several previously published methods: these include SweepFinder [aka CLR; 28], *SFselect* [37], Garud et al.’s haplotype approach using the H_12_ and H_2_/H_1_ statistics [24], Tajima’s *D* [36], and Kim and Nielsen’s *ω* [10], evolBoosting [40], and a support vector machine implemented that uses CLR and *ω* statistics (Methods). We extended *SFselect* and evolBoosting to allow for soft sweeps (Methods), and therefore refer to this classifier as *SFselect+* and evolBoosting+ in order to avoid confusion. We summarize the power of each of these approaches with the receiver operating characteristic (ROC) curve, which plots the method’s false positive rate on the *x*-axis and the true positive rate on the *y*-axis (Methods). Powerful methods that are able to detect many true positives with very few false positives will thus have a large area under the curve (AUC), while methods performing no better than random guessing are expected to have an AUC of 0.5.

We began by assessing the ability of these tests to detect selection in populations with constant population size and no population structure. First, we used test sets where the selection coefficient *α*=2*Ns* was drawn uniformly from 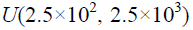, finding that S/HIC acheived had perfect accuracy (AUC=1.0; Fig S2A), and that several other methods performed nearly as well. When drawing *α* from 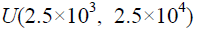, every method had near perfect accuracy (AUC>0.99) except *H*_12_ and *ω* (Fig S2B). For weaker selection 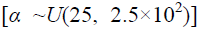 this classification task is more challenging, and the accuracies of most of the methods we tested dropped substantially. S/HIC, however, performed quite well, with an AUC of 0.9797, slightly better than evolBoosting+ (AUC=0.9702) and *SFselect+* (AUC=0.9683), and substantially better than the remaining methods (Fig S2C). Note that Garud et al.’s *H*_12_ statistic performed quite poorly in these comparisons, especially in the case of weak selection. This is likely because the fixation times of the sweeps that we simulated ranged from 0 to 0.2×*N* generations ago, and the impact of selection on haplotype homozygosity decays quite rapidly after a sweep completes [18]. Indeed, H_12_ has been shown to have good power to detect recent sweeps [24].

For the above comparisons, our classifier, evolBoosting+, and *SFselect+*, and the SVM combining CLR and *ω* were trained with the same range of selection coefficients used in these test sets. Thus, these results may inflate the performance of these methods relative to other methods, which do not require training from simulated selective sweeps. If one does not know the strength of selection, one strategy is to train a classifier using a wide range of selection coefficients so that it may be able to detect sweeps of varying strengths [37]. We therefore combined the three training sets from the three different ranges of *α* described above into a larger training set consisting of sweeps of *α* ranging from as low as 25 to as high as 25,000. This step was done not only for S/HIC, but also for *SFselect+*, evolBoosting+, and the CLR+ω SVM, and we use this approach for the remainder of the paper when using classifiers trained from constant population size data. When trained on a large range of selection coefficients, S/HIC still detected sweeps with *α* drawn from 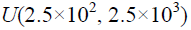 with perfect accuracy, as did *SFselect+* (Fig 2A). For stronger sweeps, we again had excellent accuracy (AUC=0.999; Fig 2B) and outperformed all other methods except *SFselect+* and SweepFinder (AUC=1.0). For weaker sweeps our method had the highest accuracy (AUC=0.9772 for S/HIC, 0.9660 for *SFselect+*, 0.9562 for evolBoosting+, and lower for other methods; Fig 2C). Thus, S/HIC can distinguish hard selective sweeps of greatly varying strengths of selection from neutrally evolving regions as well as if not better than previous methods.

**Fig. 2.**
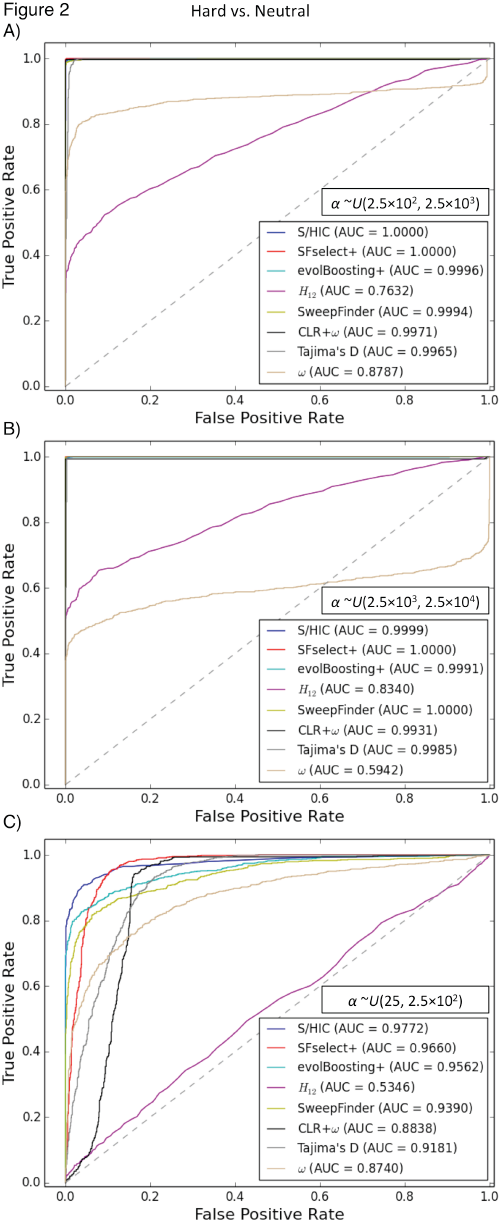
ROC curves showing the true and false positive rates of various methods/statistics when tasked with discriminating between regions containing a hard sweep and neutrally evolving regions. A) For intermediate strengths of selection (α∼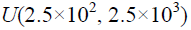)) For stronger selective sweeps 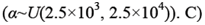. C) For weaker sweeps 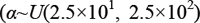. Here, and for all other ROC curves unless otherwise noted, methods that require training from simulated sweeps were trained by combining three different training sets: one where a∼*U*(2.5×10^1^, 2.5×l0^2^), one where α∼*U*(2.5×10^2^, 2.5×l0^3^), and one where α∼*U*(2.5×10^3^, 2.5×10^4^).

### S/HIC can uncover soft sweeps and distinguish them from hard sweeps

In order to uncover the targets of recent selective sweeps and also determine which mode of positive selection was responsible, one must be able to detect the signatures of soft (initial selected frequency *f* ∼U(0.05, 0.2)) as well as hard sweeps and to distinguish between them. We therefore assessed the power of each method to distinguish sets of simulated selective sweeps consisting of equal numbers of hard and soft sweeps from neutral simulations, using the same training data (for methods that require it) as for the analysis in Fig 2. For all ranges of selection coefficients, S/HIC has excellent power to distinguish hard and soft sweeps from neutrality; our AUC scores ranging from 0.9533 to 0.9862, and are higher than every other method in each scenario (Fig S3). S/HIC also distinguishes hard sweeps from soft sweeps with accuracy similar to evolBoosting+ and *SFselect+*, and these three methods perform better than each other method, except in the case of weak sweeps where *SFselect+* and *evolBoosting+* have slightly better power (AUC=0.0.7978 for S/HIC, versus 0.8066 and 0.8239, for *SFselect+* and *evolBoosting+*, respectively; Fig S4).

### Distinguishing sweeps from linked selection to narrow the target of adaptation

The goal of genomic scans for selective sweeps is not merely to quantify the extent to which positive selection impacts patterns of variation, but also to identify the targets of selection in hopes of elucidating the molecular basis of adaption. Unfortunately, hitchhiking events can skew patterns of variation across large chromosomal stretches often encompassing many loci. Furthermore, this problem not only confounds selection scans by obscuring the true target of selection, it may also lead to falsely inferred soft sweeps as a result of the soft shoulder effect [18]. Our goal in designing S/HIC was to be able to accurately distinguish among positive selection, linked selection, and neutrality, thereby addressing both of these challenging problems.

In order to assess the ability of our approach and other methods to perform this task, we repeated the test shown in Fig S3, but this time we included regions linked to selective sweeps among the set of neutral test instances. Thus, we ask how well these methods distinguish genomic windows containing the targets of selective sweeps (soft or hard) from neutrally evolving windows or windows closely linked to sweeps. Encouragingly, we find that S/HIC is able confidently distinguish windows experiencing selective sweeps from linked as well as neutrally evolving regions (Fig 3)—S/HIC achieves substantially higher accuracy than each other method (AUC=0.9593 or higher for all values of *α*, while no other method has AUC>0.91 for any *α1*). As the selection coefficient increases, S/HIC’s performance increase relative to that of other methods is particularly pronounced (Fig 3A, B), which is unsurprising because in these cases the impact of selection on variation within linked regions is much further reaching than for weak sweeps (Fig 3C).

**Fig. 3.**
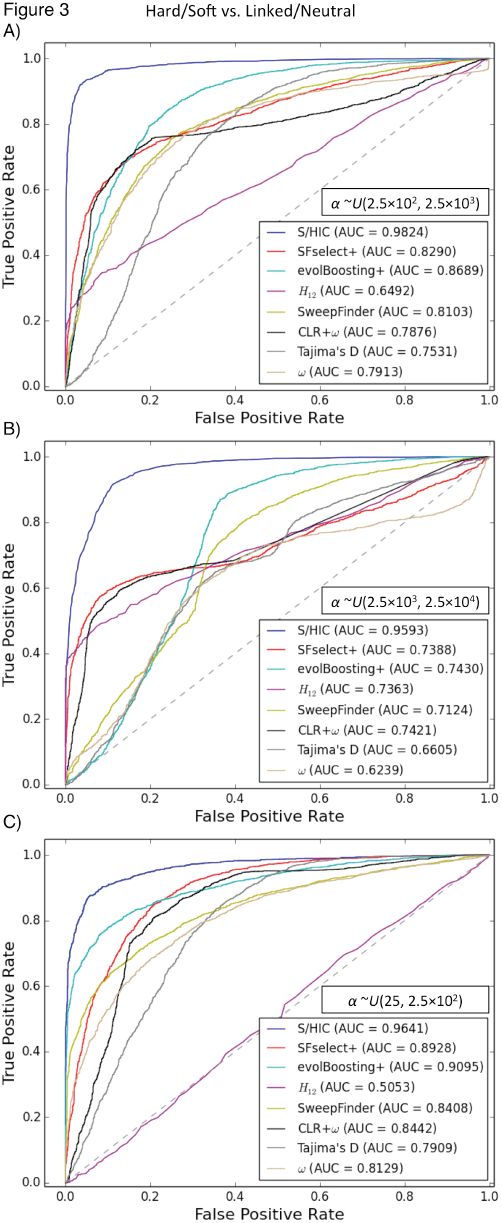
ROC curves showing the true and false positive rates of various methods/statistics when tasked with discriminating between regions containing a sweep (either hard or soft) and unselected regions (either neutral or linked to sweeps). A) For intermediate strengths of selection (α∼*U*(2.5×10^2^, 2.5×10^3^)). B) For stronger selective sweeps (α∼*U*(2.5×10^3^, 2.5×10^4^)). C) For weaker sweeps (α∼*U*(2.5×10^1^, 2.5×10^2^)).

While ROC curves provide useful information about power, a more complete view of our ability to distinguishing among hard sweeps, soft sweeps, linked selection, and neutrality can be obtained by asking how our classifier behaves at varying distances from selective sweeps. We directly compared our method’s ability to classify regions ranging from 5 subwindows upstream of a hard sweep to 5 subwindows downstream of a hard sweep to evolBoosting+ and *SFselect+* which were the top performers among all other methods we had examined. For these simulations, each subwindow had a total recombination rate 4Nr=80, corresponding to 0.2 cM per subwindow when *N*=10,000. We then counted the fraction of simulations predicted to belong to each of our five classes (hard, hard-linked, soft, soft-linked, and neutral) or the three classes used for evolBoosting+ and *SFselect+*. As shown in Fig 4, we find that when *α* ranges from 250 to 2500 and there has been a hard sweep in the central window, all three methods recover the sweep with high frequency when examining the correct window (>95% accuracy). However, as we move away from the selected site, a large number of windows are misclassified as hard sweeps by *SFSelect+* and evolBoosting+. For example, both methods misclassify nearly 50% of cases two windows away from the true sweep as hard sweeps, and most of the remaining examples as soft sweeps. In contrast, our method classifies <5% of these regions as sweeps, correctly classifying >93% of these windows as hard-linked instead. At a distance of 5 windows away from the sweep, *SFselect+* classifies the majority of windows as soft sweeps and many others as hard sweeps, and evolBoosting+ exhibits a smaller but still prominent shoulder effect (21.2% of windows classified as soft and 12.7% as hard). Meanwhile, S/HIC classifies >95% of these windows as hard-linked, and <1% as sweeps of either mode. For soft sweeps, we have the highest sensitivity in the sweep window (78.6% for S/HIC versus 75.9% for *SFselect+* and 60.8% for evolBoosting+). We also narrow the target of selection down to a smaller region, as we classify the majority of flanking windows as soft-linked, while *SFselect+* produces many soft sweep calls in these windows.

**Fig. 4.**
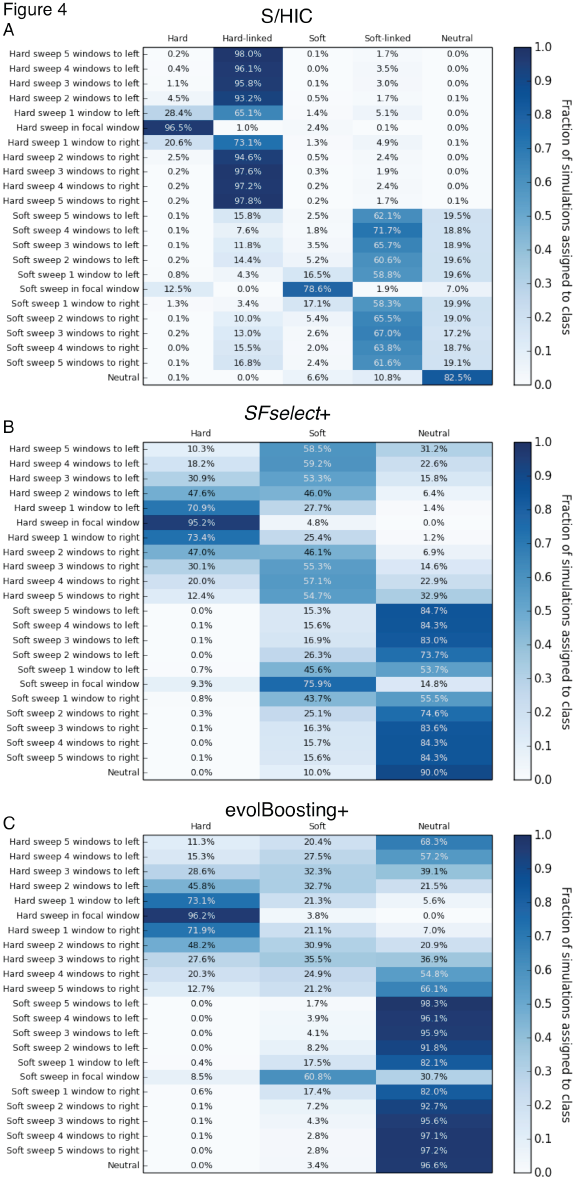
Heatmaps showing the fraction of regions at varying distances from sweeps inferred to belong to each class by S/HIC, *SFselect+*, and evolBoosting+. The location of any sweep relative to the classified window (or “Neutral” if there is no sweep) is shown on the *y*-axis, while the inferred class on the *x*-axis. Here, α∼*U*(2.5×10^2^, 2.5×l0^3^). A) Results for S/HIC. B) *SFselect+*. C) evolBoosting+.

The difference between S/HIC and these two other methods is amplified when testing these classifiers on stronger hard sweeps (*α* ranging from 2,500 to 25,000). Our classifier is better able to narrow down the selected region by classifying flanking windows as hard-linked, while *SFselect+* and evolBoosting classifies the vast majority of simulations even 5 windows away from the target of selection as hard sweeps (Fig 5). *SFselect+* and evolBoosting both have more sensitivity to detect hard sweeps when examining the correct window (>99% versus 88.8%), as S/HIC misclassifies 10.9% of these stronger sweeps as hard-linked. On the other hand, S/HIC recover 91.8% of soft sweeps versus 87.7% for *SFselect+* and 73.3% for evolBoosting+, and correctly classifies the mode of selection more often than these methods. We also misclassify relatively few regions linked to soft sweeps as sweeps themselves (∼16% when one window away, versus ∼50% for *SFselect+* and ∼20% for evolBoosting+).

**Fig. 5.**
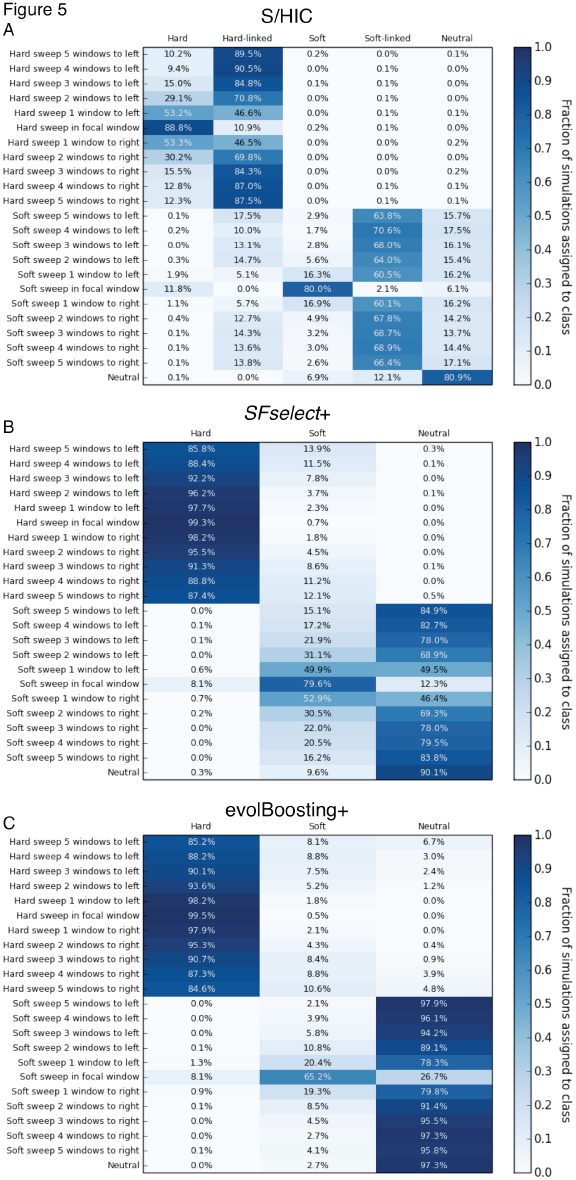
Heatmaps showing the fraction of regions at varying distances from strong sweeps inferred to belong to each class by S/HIC, *SFselect+*, and evolBoosting+. The location of any sweep relative to the classified window (or “Neutral” if there is no sweep) is shown on the y-axis, while the inferred class on the x-axis. Here, α∼*U*(2.5×10^3^, 2.5×10^4^). A) Results for S/HIC. B) *SFselect+.* C) evolBoosting+.

For weaker sweeps [*α* ∼U(25, 250)], the impact of selection on linked regions is reduced, and *SFselect+* and evolBoosting+ call fewer false sweeps in linked regions than under stronger positive selection. However, S/HIC has greater sensitivity to both hard and soft sweeps at the correct window, and also misclassifies fewer flanking regions as sweeps (Fig S5). Across the entire range of selection coefficients, S/HIC mislabeled fewer neutral simulations as sweeps than *SFselect+*, though evolBoosting+ had a slightly lower false positive rate. In summary, across all selection coefficients S/HIC has greater sensitivity than other methods to detect soft sweeps, and also for hard sweeps except when selection is very strong. Importantly, for both types of sweeps S/HIC will identify a smaller candidate region around the selective sweep than *SFselect+* or evolBoosting+. S/HIC is able to classify far fewer linked windows as selected because it has two classes for this purpose, hard-linked and soft-linked, that the other methods lack. Though *SFselect+* could be improved by incorporating these classes, it may prove difficult to determine whether a window is selected or merely linked to a sweep on the basis of its SFS alone [18], rather than examining larger scale spatial patterns of variation. evolBoosting+ fares better in this respect because it does incorporate spatial information. However, perhaps because it takes the true values of each statistic in each window rather than the relative values and also lacks “linked” classes, this method still experiences a much greater soft shoulder effect than S/HIC.

### Selection on low frequency standing variants, and ranking feature importance

Up until this point our model of selection on previously stranding variation specified an initial selected frequency, *f*, ranging from 0.05 to 0.2. However, a large fraction of soft selective sweeps may begin the sweep phase at a lower frequency [13, 16, 20]. Therefore, in order to assess how our classifier performs when soft sweeps have a lower initial selected frequency, we repeated these analyses with *f* drawn from *U*(2/2*N*, 0.05). Again, for all three ranges of the selection coefficient S/HIC has greater accuracy than any other method (Fig S6). When attempting to distinguish between hard sweeps and soft sweeps under this parameterization, performance was reduced considerably for all methods, and there was no clear winner across all strengths of selection. While S/HIC was not the top performer at this task, its AUC was within 5% of the highest score for each range of selection coefficients (Fig S7).

Next, for S/HIC and each other method that requires training, we constructed a training set in the same manner as above but allowing *f* to range from *U*(2/2*N*, 0.2), and we use this range of initial selected frequencies for all analyses presented below. While training S/HIC on these data, we ranked the importance of each feature’s contribution to our Extra-Trees classifier’s accuracy (Methods), which we list in Table S2. Generally, we find that features near the center of the window have a greater contribution, and that relative values of 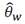 and π tend to have greater importance than other statistics.

### The impact of population size change and demographic misspecification

Non-equilibrium demographic histories have the potential to confound population genetic scans for selective sweeps [53, 54]. We therefore sought to assess the power of S/HIC and other methods to detect selection occurring in populations experiencing dramatic changes in population size. To this end we trained and tested our classifiers under four demographic scenarios (Table S1): two simple population bottlenecks of varying severity (one of which models European *Drosophila*), a model of recent exponential population size growth, and finally a more complex model that describes out-of-Africa populations of humans.

#### Human demographic models

We examined the performance of S/HIC under demographic models that were recently estimated for African and European human populations by Tennessen et al. [44]. The African model from Tennessen et al. consists of recent exponential growth in population size. The European model from Tennessen et al. (Table S1) includes recurrent population contractions followed by first slow and then accelerated population growth. Performance of these models is shown in Fig S8, from which it can be seen that S/HIC has the highest accuracy of all methods that we examined. For these two scenarios both training and testing data were drawn from the same demographic model.

A more pessimistic scenario is one where the true demographic history of the population is not known, and therefore misspecified during training. Most demographic events should impact patterns of variation genome-wide rather than smaller regions (but see refs. [55, 56]). Thus, approaches that search for spatial patterns of polymorphism consistent with selective sweeps may be more robust to demographic misspecification than methods examining local levels of variation only (as demonstrated by ref. [28]). To test this, we trained S/HIC and other classifiers on equilibrium datasets, and measured their accuracy on test data simulated under the non-equilibrium demographic models described above. In Fig S9A we show the power of these classifiers to detect selective sweeps occurring under the African model of recent exponential growth. Under this scenario, with 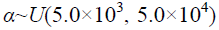 (equivalent to 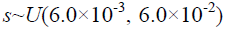 with *N*=424,000), S/HIC achieves an AUC of 0.8122, while the next-highest performing method is evolBoosting+ (AUC=0.7567). Similarly, we perform better than other methods when searching for stronger selection (α ranging from 5.0×10^4^ to 5.0×10^5^; AUC=0.9844 versus <0.92 for all others; Fig S9B).

Note that the simple summary statistic methods *ω* and Tajima’s *D* have some power to detect selection even under non-equilibrium demography (Fig S8). However, this result is probably quite optimistic: the ROC curve is generated by repeatedly adjusting the critical threshold and measuring true and false positive rates. In practice, a single critical threshold may be chosen to identify putative sweeps. If this critical value is chosen based on values of the statistic generated under the incorrect demographic model, then the false positive rate may be quite high. For example, Nielsen et al. [28] showed that when a threshold for Tajima’s *D* is selected based on simulations under equilibrium, 100% of neutral simulations under a population growth model exceed this threshold. In other words, the ROC curve is useful for illustrating a method’s potential power if an appropriate threshold is selected, but this may not always be the case in practice.

A more informative approach to evaluating our power may thus be to examine the fraction of regions including sweeps, linked to sweeps at various recombination distances, or evolving neutrally, that were assigned to each class (as done in Figs 4-5 for constant population size). We show this in Fig S10, which better illustrates S/HIC’s power and robustness to unknown demographic history. Overall, S/HIC has roughly similar sensitivity to selection as *SFselect+* and evolBoosting+. For example, with *α* 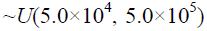, we recover 98.3% of hard sweeps versus 99.7% for *SFselect+* and 99.3% for evolBoosting+ (Fig S10D-F), though these three methods misclassify many of hard sweeps as soft (48.5%, 26.8%, and 30.6%, respectively). For soft sweeps, with 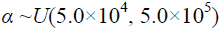, S/HIC classifies 84.5% of examples correctly, and an additional 7.9% as hard, versus 83.0% as soft and 12.3% as hard for *SFselect+*, and 77.6% as soft and 8.4% as hard for evolBoosting+. When examining windows linked to selective sweeps, both *SFselect+* and evolBoosting+ incorrectly classify large fractions of instances as hard or soft sweeps (especially for stronger selection coefficients), while S/HIC classifies most of these as hard-linked or soft-linked (or neutral in the case of weak selection)— indeed our method classifies very few linked regions as selective sweeps.

In the context of scans for positive selection, the primary concern with non-equilibrium demography is that it will produce a large number of false selective sweep calls. Indeed, when trained on an equilibrium demographic history and tested on the exponential growth model, *SFselect+* classifies roughly one-fifth of all neutral loci as having experienced recent positive selection; for evolBoosting+ the false positive rate is ∼15%. In stark contrast, S/HIC does not seem to be greatly affected by this problem: we classify only ∼7% of neutrally evolving regions as sweeps. Thus, while these three methods all have comparable sensitivity to sweeps in this scenario, S/HIC has superior specificity: *SFselect+* and evolBoosting+ classify a large fraction of unselected regions (including both neutral and sweep-linked regions) as sweeps, whereas S/HIC has a low false positive rate. For the curious reader, we present S/HIC’s feature rankings for classifiers trained on both human demographic histories in Table S2.

Next, we examined the impact of demographic misspecification on power to detect selection occurring under Tennessen et al.’s model of the population size history of Europeans following their migration out of Africa [44] but having trained S/HIC under the standard neutral model. This demographic history presents an even greater challenge for identifying positive selection than the African model, as it is characterized by two population contractions followed by exponential growth, and then a more recent phase of faster population growth (Methods). For this scenario, a single range of selection coefficients was used: 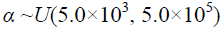. Here, we find that, perhaps unsurprisingly, the performance of most methods is lower than in the African scenario. However, S/HIC once again appears substantially more robust to misspecification of the demographic model than other methods (AUC=0.8127 versus 0.7250 for evolBoosting+, and ∼0.6 or less for all other methods; Fig 6).

**Fig. 6.**
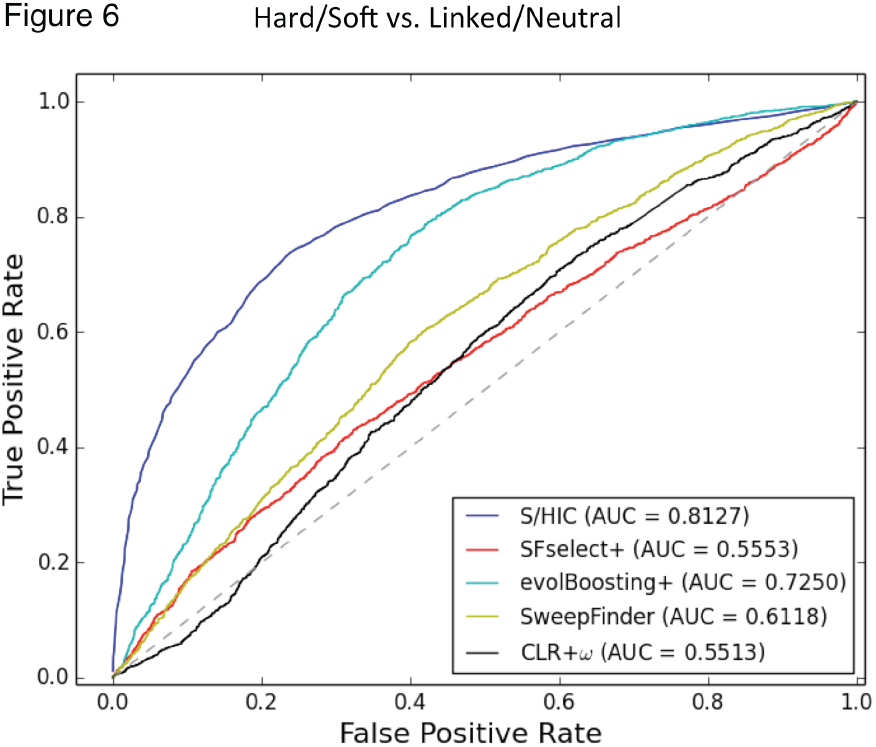
ROC curves showing the true and false positive rates of various methods/statistics when tasked with discriminating between regions containing a sweep (either hard or soft) and unselected regions (either neutral or linked to sweeps) when testing on simulations with Tennessen et al.’s European demographic model. Here, α∼*U*(5×10^3^, 5×10^5^), and the methods that require training from simulated sweeps were trained from the same simulations with equilibrium demography as used for Figs 2-5. Note that Tajima’s *D* and Kim and Nielsen’s *ω* were omitted from this figure, as we simply used the values of these statistics to generate ROC curves without respect to any demographic model.

Next, we examined the proportion of windows at various distances from sweeps that are assigned to each class under this scenario of demographic misspecification. We find that while S/HIC classifies hard sweeps with lower sensitivity than under constant population size scenario (56.0% and 19.1% of test examples are classified as hard and soft, respectively), relatively few linked windows are classified as sweeps (Fig 7A). For soft sweeps S/HIC fares less well (20.7% of windows are correctly classified, and 34.7% classified as hard sweeps), though again relatively few false positives are produced in linked or neutral regions. In contrast, evolBoosting+ classifies the majority of windows, selected or otherwise, as soft sweeps (Fig 7C): 68.5% of hard sweeps and 55.0% of neutral regions are misclassified as soft. For SFselect+ this problem in exacerbated: 68.6% of hard sweeps and 95.3% of neutral windows are classified as soft sweeps. Thus, under this scenario of demographic misspecification, S/HIC is the only method we examine which can discriminate between positively selected and unselected portions of the genome effectively.

**Fig. 7.**
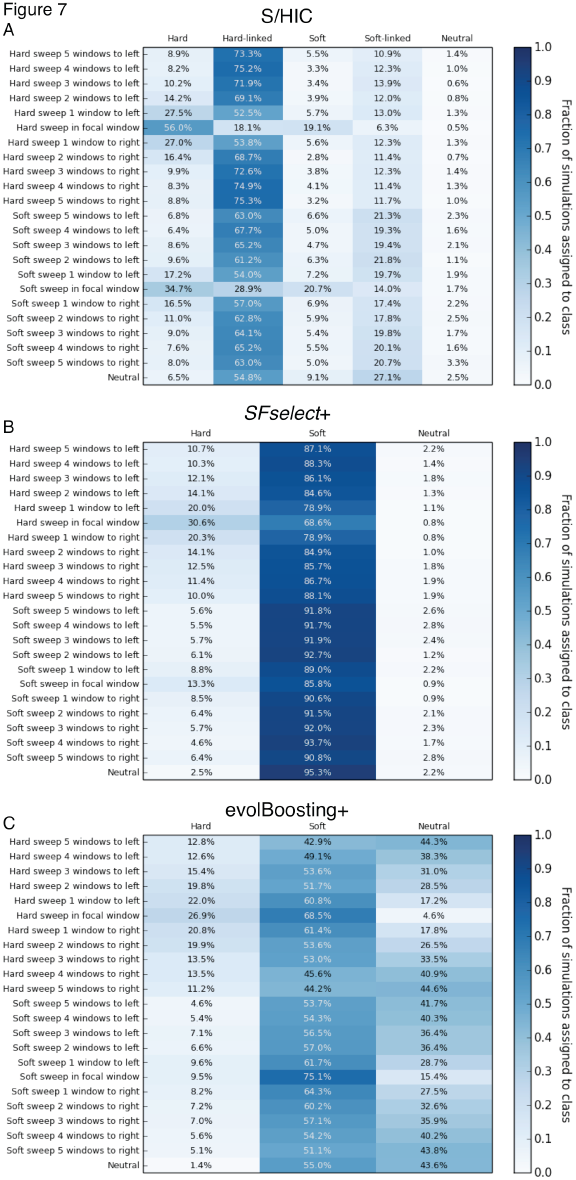
Heatmaps showing the fraction of regions simulated under Tennessen et al.’s European demographic model located at varying distances from sweeps inferred to belong to each class by S/HIC, *SFselect+*, and evolBoosting+. The location of any sweep relative to the classified window (or “Neutral” if there is no sweep) is shown on the *y*-axis, while the inferred class on the *x*-axis. Here, α∼*U*(5×10^3^, 5×10^5^). These three classifiers were trained from simulations with equilibrium demography. A) Results for S/HIC. B) *SFselect+*. C) evolBoosting+.

#### Bottleneck models

Next we sought to examine accuracy on a 3-epoch population bottleneck in which the population contracts to a much smaller size and then later recovers to its original size. We first examined a model similar to that from Thornton and Andolfatto [47] for European *Drosophila* populations, but with a less severe contraction in population size (Methods; Table S1). We assessed each method’s ability to detect sweeps with three different fixation times: the immediate past, in the middle of the bottleneck, and immediately prior to the population contraction. When methods that require training were trained under the correct demographic model, S/HIC was better able to discriminate between sweeps and unselected windows than any other method (Fig S11). This difference grew more pronounced with increasing time since the sweep: for instance, S/HIC had an AUC of 0.9458 for the oldest fixation time considered, while the next most accurate method was evolBoosting+ with an AUC of 0.8023 (Fig S11C). Furthermore, when all methods were trained from simulations under constant population size (i.e. misspecified), their performance tended to decrease, sometimes considerably, while S/HIC exhibited no drop in accuracy (Fig S11D-F).

We then tested our set of tools on a bottleneck model with the same parameterization as Thornton and Andolfatto’s model estimated in *D. melanogaster* [47]. This bottleneck is 10-fold more severe than the model tested above. In this model, the population size decreases to just 2.9% of its original size during the bottleneck. Detecting positive selection in the context of this population size history is far more challenging, and the performance of every method suffers considerably (Fig S12). Still, S/HIC is once again the top performing method for both recent fixations and the oldest fixation time we examined, though SFselect+ is slightly more powerful for the intermediate fixation time (AUC=0.6902 verus 0.6750). When trained on equilibrium demography S/HIC experiences no drop in accuracy for older sweeps, in contrast to the other methods (e.g. on very old sweeps SFselect+ performs worse than a random classifier). Interestingly, S/HIC does show a noticeable drop in AUC (from 0.9182 to 0.7817) when trained on the wrong model, but still has the highest accuracy in this case (Fig S12A, D). Thus under each demographic model we examined, S/HIC exhibits sensitivity to selective sweeps comparable to other top-performing methods (though it occasionally struggles to correctly infer the mode of selection). More importantly, S/HIC avoids the enormous false positive rates that may plague other methods. Taken together, the above results lend credence to the idea that spatial patterns of variation will be more robust to non-equilibrium demography, and far less impaired by misspecification of the demographic model.

### Identifying selective sweeps in a human population sample with European ancestry

The results from simulated data described above suggest that our method has the potential to identify selective sweeps and distinguish them from linked selection and neutrality with excellent accuracy. In order to demonstrate our method’s practical utility, we used it to perform a scan for positive selection in humans. In particular, we searched the 1000 Genomes Project’s CEU population sample (European individuals from Utah) for selective sweeps occurring after the migration out of Africa. We focused this search on chromosome 18, where several putative selective sweeps have been identified in Europeans [57]. The steps we took to train our classifier and filter the 1000 Genomes data prior to conducting our scan are described in the Methods.

In total, we examined 344 windows, each 200 kb in length. We classified 34 windows (9.9%) as centered around a hard sweep, 22 (6.4%) as linked to a hard sweep, 48 (14.0%) as centered around a soft sweep, 89 (25.9%) as linked to a soft sweep, and 151 (43.9%) as neutral. Surprisingly, we infer that over 56% of windows lie within regions whose patterns of variation are affected by sweeps either within the window or in linked regions. This may imply that, given the genomic landscape of recombination in humans, even if selective events are somewhat rare [58], they may nonetheless impact variation across large stretches of the genome. However, we cannot firmly draw this conclusion given the difficulty of distinguishing between linked selection and neutrality under the European demographic model (Fig 7).

Encouragingly, our scan recovered 4 of the 5 putative sweeps on chromosome 18 in Europeans identified by Williamson et al. [57] using SweepFinder. These include *CCDC178* (which we classify as a hard sweep), *DTNA* (which we classify as soft), *CCDC102B* (hard), and the region spanning portions of *CD226* and *RTTN* (hard). In each of these loci, the windows that we predicted to contain the sweep overlapped regions of elevated composite likelihood ratio (CLR) values from SweepFinder [visualized using data from 59]. Although the CLR statistic is not completely orthogonal to the summary statistics we examine to perform our classifications, the close overlap that we observe between these two methods underscores our ability to precisely detect the targets of recent positive selection. We also identify a novel candidate hard sweep within *L3MBTL4*, an apparent tumor suppressor gene that is often mutated, downregulated, or deleted in breast tumors [60]. As shown in Fig. 8, π, Tajima’s *D*, *Z_nS_*, *ω*, and the CLR statistic all show patterns strongly suggestive of a selective sweep within this gene. The complete set of coordinates of putative sweeps from this scan is listed in Table S4.

**Fig. 8.**
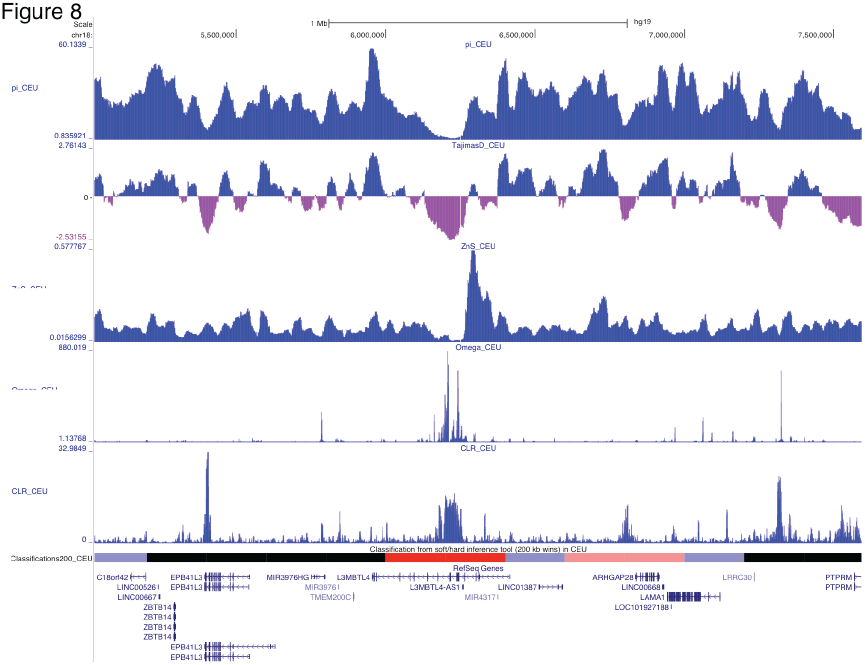
Browser screenshot showing patterns of variation around a putative selective sweep in Europeans within *L3MBTL4* on chr18. Values of *π*, Tajima’s *D*, Kelley’s *Z_nS_*, and Nielsen et al’s composite likelihood ratio, all from Pybus et al. [59], are shown. Beneath these statistics we show the classifications from S/HIC (red: hard sweep; faded red: hard-linked; blue: soft sweep; faded blue: soft-linked; black: neutral). This image was generated using the UCSC Genome Browser (http://genome.ucsc.edu).

Next, we asked whether S/HIC recovered evidence of positive selection on the *LCT* (lactase) locus. Previous studies have found evidence for very recent and strong selection on this gene in the form population differentiation and long-range haplotype homozygosity [61–63]. Moreover, several variants in this region are associated with lactase persistence. Nielsen et al.’s CLR has also identified this region [28], but not consistently: Williamson et al.’s [57] CLR scan did not detect a sweep at this locus, nor does a recent scan using the 1000 Genomes data [data from 59]. This may be expected, as the selection on lactase persistence alleles appears to have not yet produced completed sweeps. Overall, there is very strong evidence of recent and perhaps ongoing selection for lactase persistence in human populations relying on diary for nutrition.

Like the SweepFinder CLR, S/HIC in its current form is also designed to detect completed sweeps. Nonetheless, we applied S/HIC to a 4 Mb region on chromosome 2 spanning *LCT* and neighboring loci. Consistent with previous studies, we find evidence of a selective sweep in a region upstream of *LCT* (Fig S13) that contains a mutation associated with lactase persistence in Europeans [64], though unlike Peter et al. [25], we classify this sweep as soft. We also find evidence of a hard selective sweep upstream of *LCT*, suggesting that there may be additional targets of selection in this region of chromosome 2. Consistent with this is the observation that our candidate window also overlaps a region identified by Green et al. [65] as having an excess of derived alleles in the human genome relative to the number observed in Neanderthal.

The computational speed of this scan is largely governed by the amount of time spent calculating summary statistics to generate the feature vectors (as well as simulating training data, if absent), as the training and classification tasks are relatively inexpensive (typically requiring several minutes for the former and only seconds for the later). The approximate runtime for calculating our set of summary statistics within the 4 Mb region encompassing *LCT* is ∼30 minutes (using code from https://github.com/kern-lab/shIC). Thus, if a compute cluster is available, one can subdivide the genome into segments of this size and perform these calculations in parallel, and classify every window in the human genome in under an hour.

## DISCUSSION

Detecting the genetic targets of recent adaptation and the mode of positive selection acting on them—selection on *de novo* mutations versus previously standing variants—remains an important challenge in population genetics. The majority of efforts to this end have relied on population genetic summary statistics designed to uncover loci where patterns of allele frequency [e.g. 8, 36, 66] or linkage disequilibrium [e.g. 9, 10] depart from the neutral expectation. Recently, powerful machine learning techniques have begun to be applied to this problem, showing great promise [18, 37, 39, 40, 43]. Here we have adopted a machine learning approach to develop S/HIC, a method designed to not only uncover selective sweeps, but to distinguish them from regions linked to sweeps as well as neutrally evolving regions, and to identify the mode of selection. This is achieved by examining spatial patterns of a variety of population genetic summary statistics that capture different facets of variation across a large-scale genomic region. Currently, this method examines the values of nine statistics across eleven different windows in infer the mode of evolution in the central window—this makes for a total of 99 different values considered by the classifier. By leveraging all of this information jointly, our Extra-Trees classifier is able to detect selection with accuracy unattainable by methods examining a single statistic, underscoring the potential of the machine learning paradigm for population genetic inference. Indeed, on simulated datasets with constant population size, S/HIC has power matching or exceeding previous methods when linked selection is not considered (i.e. the sweep site is known *a priori*), and vastly outperforms them under the more realistic scenario where positive selection must be distinguished from linked selection as well as neutrality.

We argue that the task of discriminating between the targets of positive selection and linked but unselected regions is an extremely important and underappreciated problem that must be solved if we hope to identify the genetic underpinnings of recent adaptation in practice. This is especially so in organisms where the impact of positive selection is pervasive, and therefore much of the genome may be linked to recent selective sweeps [e.g. 67]. A method that can discriminate between sweeps and linked selection would have three important benefits. First, it will reduce the number of spurious sweep calls in flanking regions, thereby mitigating the soft shoulder problem [18]. Second, such a method would have the potential to narrow down the candidate genomic region of adaptation. Third, such a method would be able to find those regions *least* affected by linked selection, which themselves might act as excellent neutral proxies for inference into demography or mutation. We have shown that S/HIC is able to distinguish among selection, linked selection, and neutrality with remarkable power, granting it the ability to localize selective sweeps with unrivaled accuracy and precision, demonstrating its practical utility.

While S/HIC performs favorably to other approaches under the ideal scenario where the true demographic history of the population is known, in practice this may not always be the case. However, because our method relies on spatial patterns of variation, we are especially robust to demography: if the demographic model is misspecified, the disparity in accuracy between S/HIC and other methods is even more dramatic. For example, if we train S/HIC with simulated datasets with constant population size, but test it on simulated population samples experiencing recent exponential growth (e.g. the African model from ref. [44]), we still identify sweeps with impressive accuracy, and vastly outperform other methods. We also tested S/HIC on a more challenging model with two population contractions followed by slow exponential growth, and more recent accelerated growth (the European model from ref. [44]), obtaining qualitatively similar results. S/HIC therefore seems well suited for inference on populations with unknown demographic histories, though in such scenarios power could perhaps be improved by quickly fitting a relatively simple non-equilibrium demographic model prior to training. Even if oversimplified, simulations under such a model might better approximate patterns of variation around sweeps and within unselected regions than simulations under equilibrium, though we have not explored this possibility here.

Though S/HIC performs far better than other tests for selection when tested on non-equilibrium populations, power for all methods is far lower than under constant population size, even if the demographic model is properly specified during training. Similar results are obtained under a severe population bottleneck. The reason for this is somewhat disconcerting: under these demographic models, the impact of selective sweeps on genetic diversity is blunted, making it far more difficult for any method to identify selection and discriminate between hard and soft sweeps. This underscores a problem that could prove especially difficult to overcome. That is, for some demographic histories all but the strongest selective sweeps may produce almost no impact on diversity for selection scans to exploit.

A second and related confounding effect of misspecified demography is that following population contraction and recovery/expansion, much of the genome may depart from the neutral expectation, even if selective sweeps are rare. By examining the relative levels of various summaries of variation across a large region, rather than the actual values of these statistics, we are quite robust to this problem (Fig 7 and Fig S10). In other words, while non-equilibrium demography may reduce S/HIC’s sensitivity to selection and its ability to discriminate between hard and soft sweeps, we still classify relatively few neutral or even linked regions as selected. Thus, although inferring the mode of positive selection with high confidence may remain extremely difficult in some populations, our method appears to be particularly well suited for detecting selection in populations with non-equilibrium demographic histories whose parameters are uncertain. Indeed, applying our approach to chromosome 18 in a European human population, we detect most of the putative sweeps previously reported by Williamson et al. [57].

An additional advantage of machine learning approaches such as ours is the relative ease with which the classifier can be extended to incorporate more features, potentially adding information complementary to current features that could further improve classification power. For example, our examination of linkage disequilibrium is limited to within each subwindow; including features measuring the degree of LD between subwindows could also add valuable information. In addition, we could add statistics currently omitted which capture patterns of genealogical tree imbalance (e.g. the maximum frequency of derived alleles [68]), or star-like sub-trees within genealogies (e.g. *iHS* [42], *nS*_L_ [23]), both symptoms of various types of positive selection. Indeed, all tests for selective sweeps can be seen as methods to detect the distortions in the shapes of genealogies surrounding selected sites. Thus, if one could directly examine the ancestral recombination graph (ARG) surrounding a focal region, more powerful inference could be possible. It is now possible to estimate ARGs from sequence data [69], and summaries of these estimated trees could be incorporated as features to identify sweeps and classify their mode. These are just some of a multitude of possible features that one can use to make inferences about natural selection. The success of S/HIC, evolBoosting [40], and *SFselect* [37] in our tests relative to more conventional methods shows that machine learning approaches leveraging many different types of information have the potential to make far more powerful inferences than methods relying on an individual statistic.

In summary, we have devised a machine learning-based scan for positive selection that possesses not only unparalleled accuracy, but is also exceptionally robust to the non-equilibrium demographic models we have examined here. This finding is extremely encouraging, though we can’t be certain it generalizes to every possible demographic scenario. Adjustments to the feature space can easily be made to better suit a particular study population. For example, if haplotypic phase is unknown, one can replace measures of gametic LD with zygotic LD. Additional classes could also be incorporated into the classifier (e.g. “partial” or incomplete sweeps, balancing selection, or background selection), as long as they can be simulated to generate training data. Thus, our approach is practical and flexible. As additional population genetic summary statistics and tests for selection are devised, they can be incorporated into our feature space, thereby strengthening an already powerful method which has the potential to illuminate the impact of selection on genomic variation with unprecedented detail.

## ACKNOWLEDGEMENTS

We thank Matthew Hahn and Adam Siepel for comments on the manuscript, and Yun Song for helpful conversations during this effort.

---

**Fig. S1.**
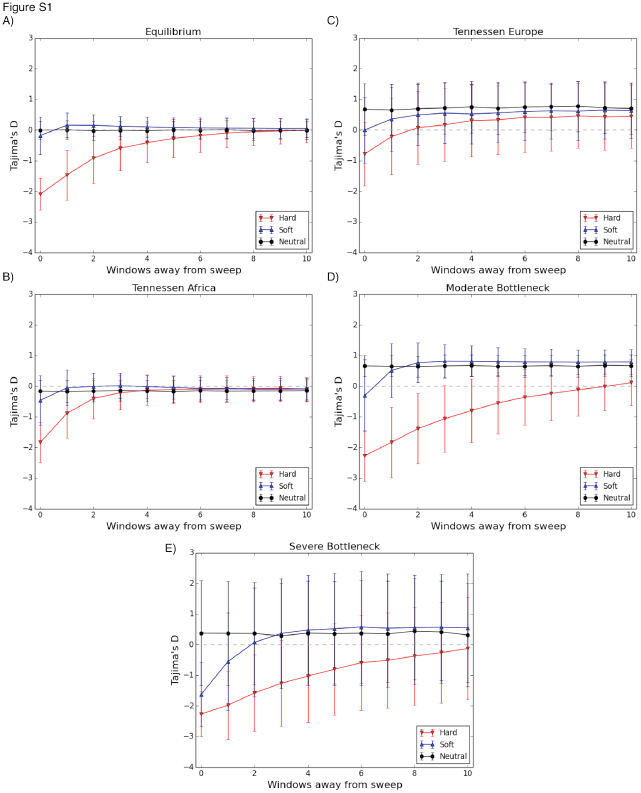
Means and standard deviations of Tajima’s *D* at increasing distances from a selective sweep, and in neutrally evolving windows. The sweep occurs in window 0. A) Values of Tajima’s *D* in 11 subwindows for the constant population size scenario, with *α* drawn from *U*(250, 2500). B) The African demographic model, with *α* drawn from *U*(5.0×10^3^, 5.0×10^4^). C) The European demographic model. D) The less severe bottleneck model (reduction to 29% of original size), with sweeps completing immediately prior to sampling. E) The more severe Thornton and Andolfatto [47] model (reduction to 2.9% of original size), with sweeps completing immediately prior to sampling.

**Fig. S2.**
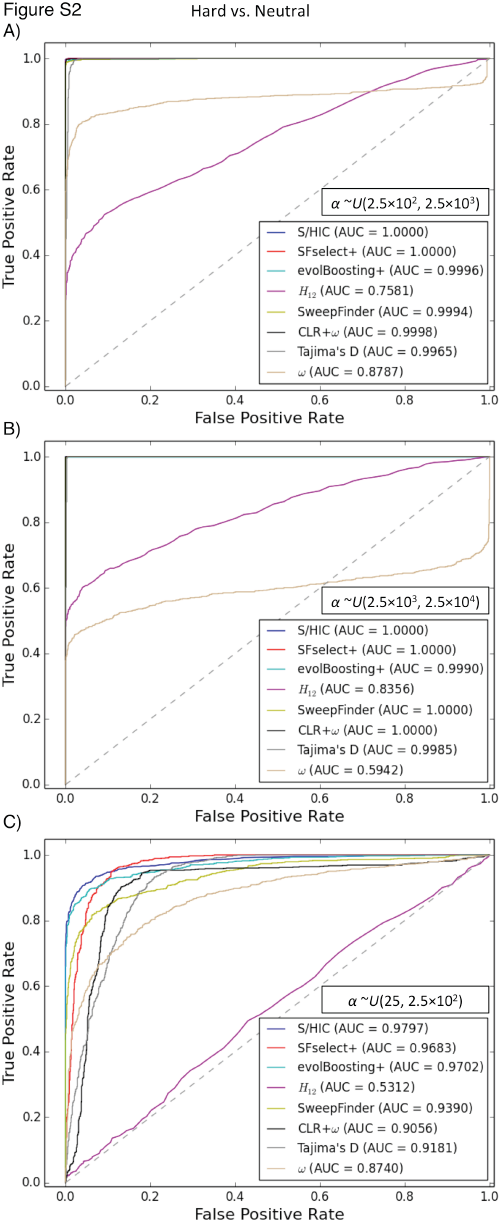
ROC curves showing the true and false positive rates of various methods/statistics when tasked with discriminating between regions containing a hard sweep and neutrally evolving regions. A) For intermediate strengths of selection (α∼*U*(2.5×10^2^, 2.5×10^3^)). B) For stronger selective sweeps (α∼*U*(2.5×10^3^, 2.5×10^4^)). C) For weaker sweeps (α∼*U*(2.5×10^1^, 2.5×10^2^)). Here, the methods that require training from simulated sweeps were trained from a set having the same distribution of selection coefficients as the test set.

**Fig. S3.**
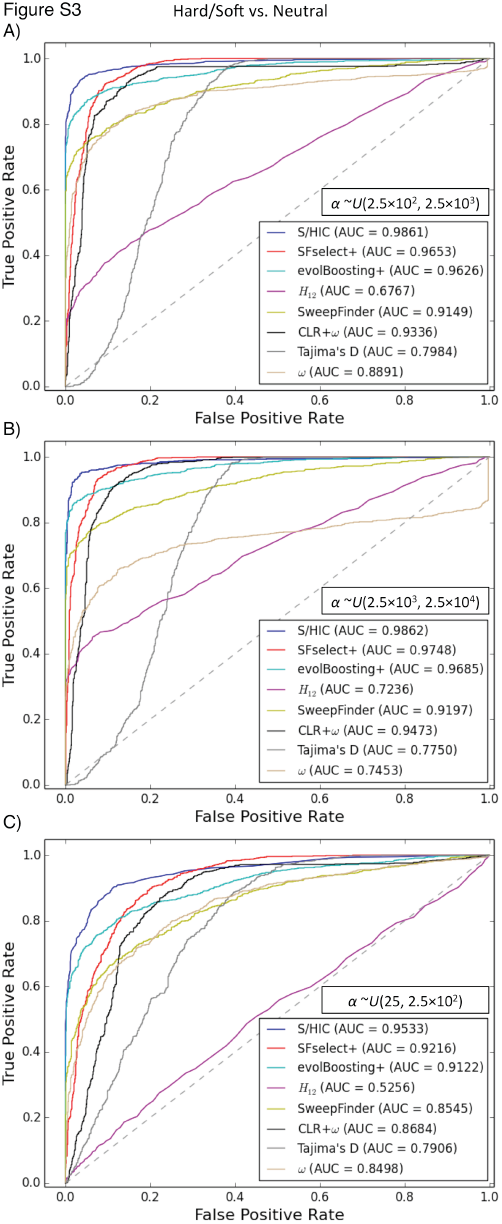
ROC curves showing the true and false positive rates of various methods/statistics when tasked with discriminating between regions containing a sweep (either hard or soft) and neutrally evolving regions. A) For intermediate strengths of selection (α∼*U*(2.5×10^2^, 2.5×10^3^)). B) For stronger selective sweeps (α∼*U*(2.5×10^3^, 2.5×10^4^)). C) For weaker sweeps (α∼*U*(2.5×10^1^, 2.5×10^2^)).

**Fig. S4.**
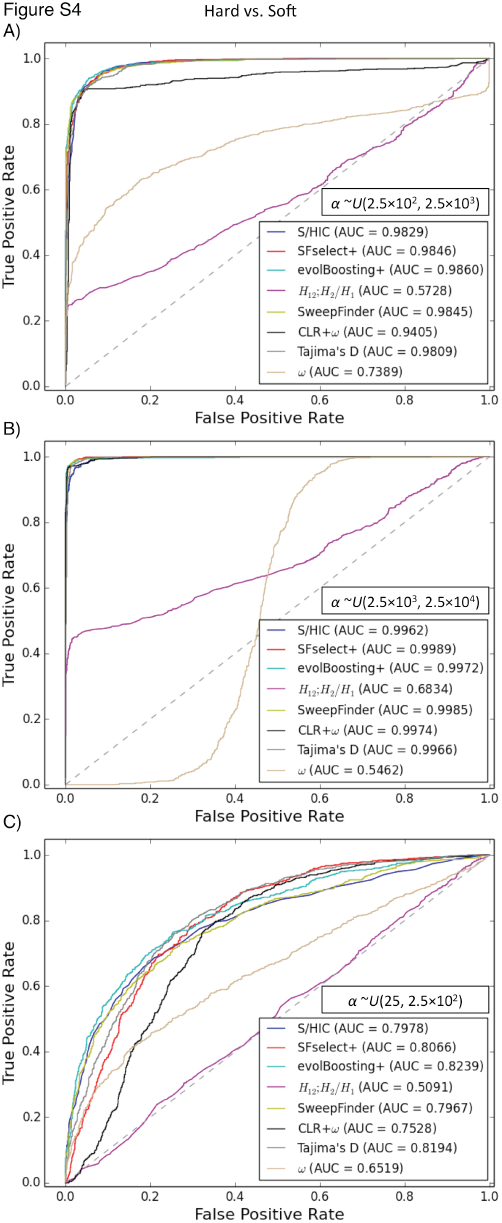
ROC curves showing the true and false positive rates of various methods/statistics when tasked with discriminating between hard and soft sweeps. A) For intermediate strengths of selection (α∼*U*(2.5×10^2^, 2.5×10^3^)). B) For stronger selective sweeps (α∼*U*(2.5×10^3^, 2.5×10^4^)). C) For weaker sweeps (α∼*U*(2.5×10^1^, 2.5×10^2^)).

**Fig. S5.**
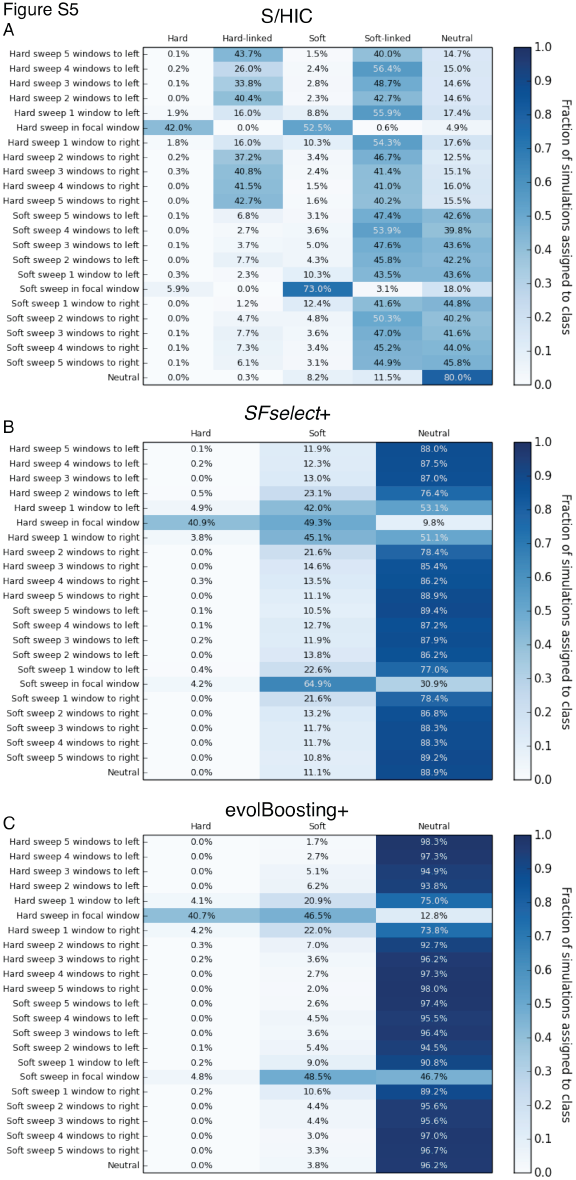
Heatmaps showing the fraction of regions at varying distances from weak sweeps inferred to belong to each class by S/HIC, *SFselect+*, and evolBoosting+. The location of any sweep relative to the classified window (or “Neutral” if there is no sweep) is shown on the *y*-axis, while the inferred class on the *x*-axis. Here, α∼*U*(2.5×10^1^, 2.5×10^2^).

**Fig. S6.**
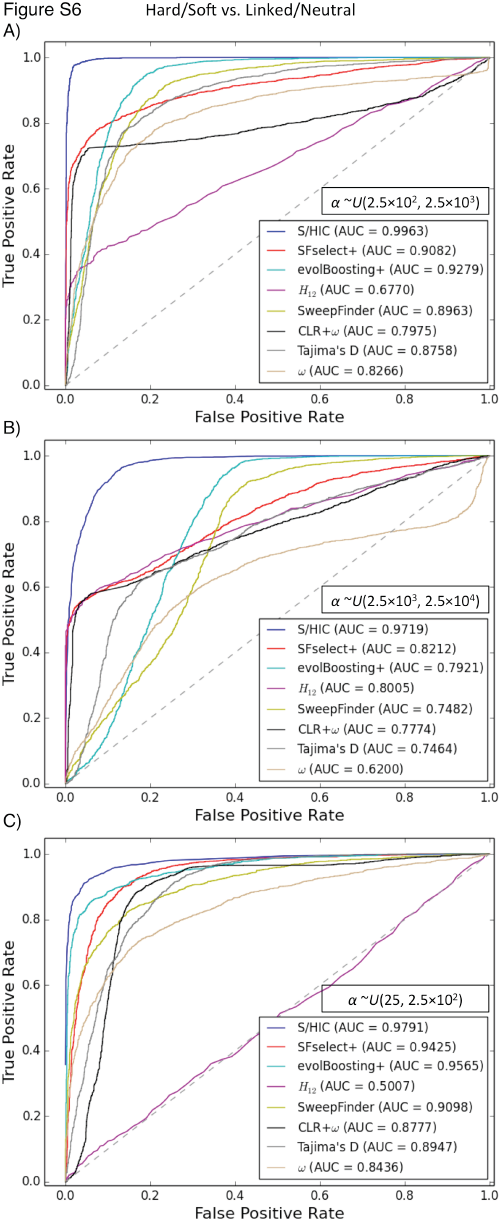
ROC curves showing the true and false positive rates of various methods/statistics when tasked with discriminating between regions containing a sweep (either hard or soft) and unselected regions (either neutral or linked to sweeps). A) For intermediate strengths of selection (α∼*U*(2.5×10^2^, 2.5×10^3^)). B) For stronger selective sweeps (α∼*U*(2.5×10^3^, 2.5×10^4^)). C) For weaker sweeps (α∼*U*(2.5×10^1^, 2.5×10^2^)). For the soft sweep training and test examples used to generate these plots, *f* was drawn from ∼*U*(2/2*N*, 0.05).

**Fig. S7.**
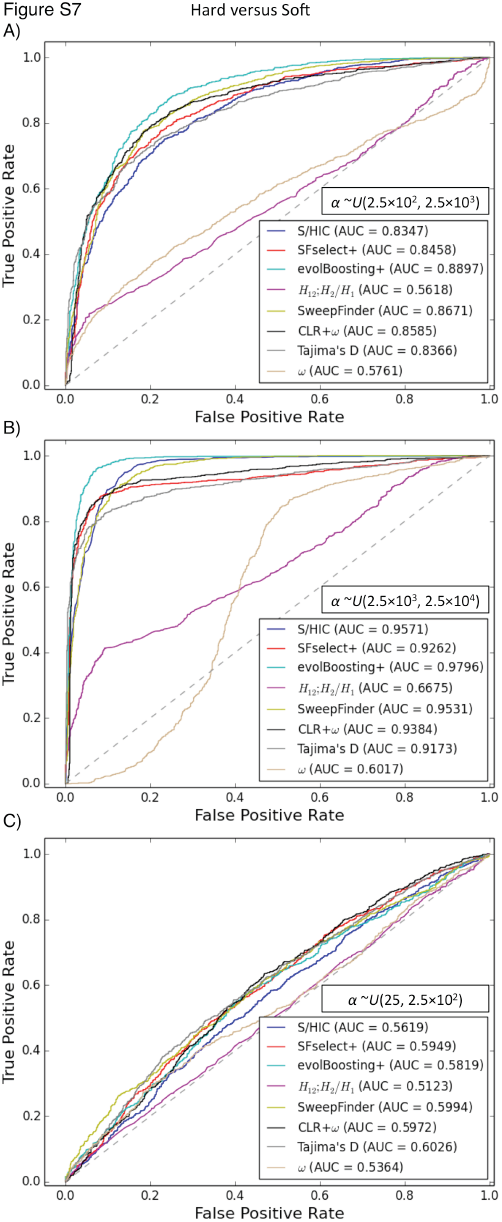
ROC curves showing the true and false positive rates of various methods/statistics when tasked with discriminating between hard and soft sweeps. A) For intermediate strengths of selection (α∼*U*(2.5×10^2^, 2.5×10^3^)). B) For stronger selective sweeps (α∼*U*(2.5×10^3^, 2.5×10^4^)). C) For weaker sweeps (α∼*U*(2.5×10^1^, 2.5×10^2^)). For the soft sweep training and test examples used to generate these plots, *f* was drawn from ∼*U*(2/2*N*, 0.05).

**Fig. S8.**
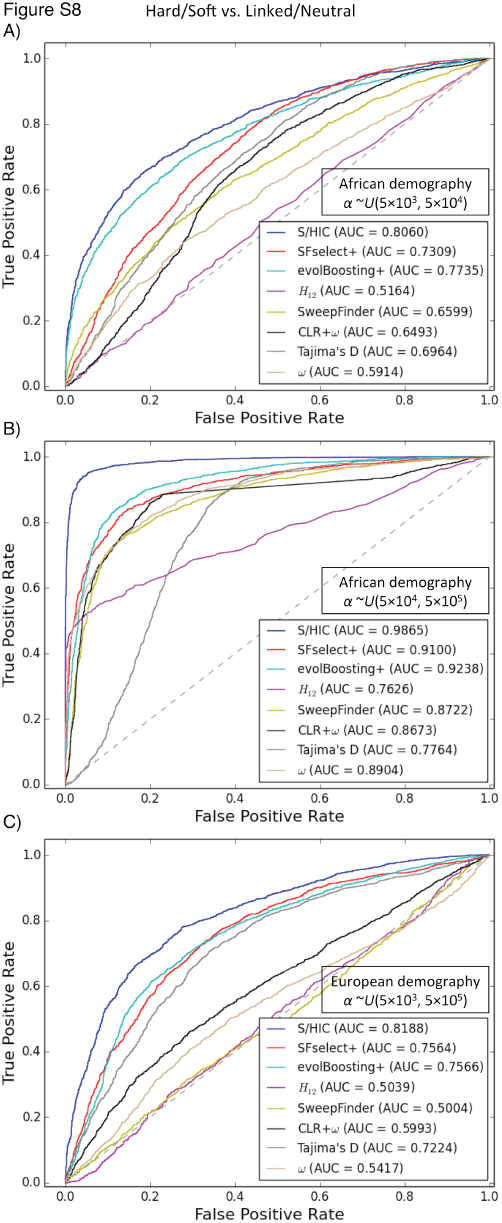
ROC curves showing the true and false positive rates of various methods/statistics when tasked with discriminating between regions containing a sweep (either hard or soft) and unselected regions (either neutral or linked to sweeps) when testing on non-equilibrium demography. Here, the methods that require training from simulated sweeps were trained from the same demographic model used for testing. A) Testing on the African demographic model, with α∼*U*(5×10^3^, 5×10^4^). B) The African demographic model, with α∼*U*(5×10^4^, 5×10^5^). C) The European demographic model, with α∼*U*(5×10^3^, 5×10^5^).

**Fig. S9.**
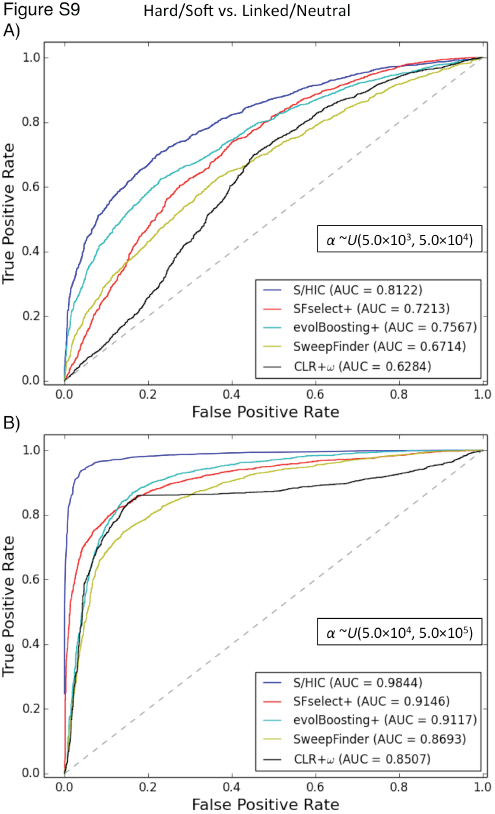
ROC curves showing the true and false positive rates of various methods/statistics when tasked with discriminating between regions containing a sweep (either hard or soft) and unselected regions (either neutral or linked to sweeps) when training with equilibrium demography but testing on non-equilibrium demography. Here, the methods that require training from simulated sweeps were trained from the same simulations with equilibrium demography as used for Figs 2-5. A) Testing on the African demographic model, with α∼*U*(5×10, 5×10^4^). B) The African demographic model, with α∼*U*(5×10^4^, 2.5×10^5^). Note that Tajima’s *D* and Kim and Nielsen’s *ω* were omitted from this figure, as we simply used the values of these statistics to generate ROC curves without respect to any demographic model.

**Fig. S10.**
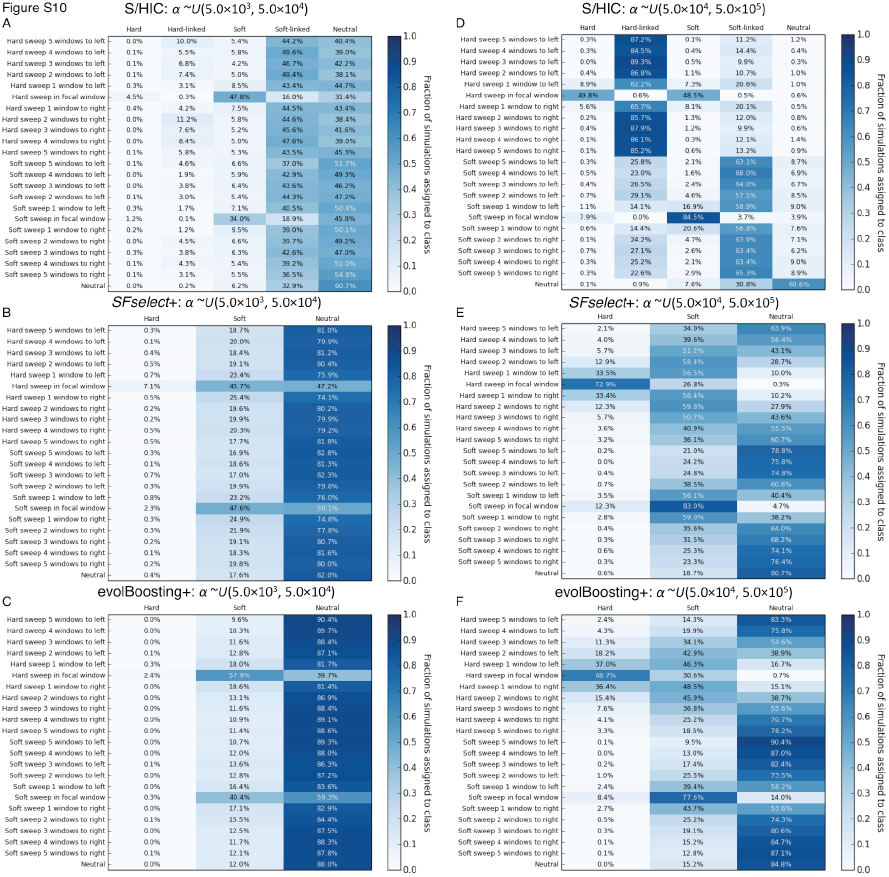
Heatmaps showing the fraction of regions simulated under Tennessen et al.’s African demographic model located at varying distances from sweeps inferred to belong to each class by S/HIC, *SFselect+*, and evolBoosting+. The location of any sweep relative to the classified window (or “Neutral” if there is no sweep) is shown on the y-axis, while the inferred class on the *x*-axis. For panels A-C, α∼*U*(5×10^3^, 5×10^4^), and for D-F α∼*U*(5×10^4^, 5×10^5^). These three classifiers were trained from simulations with equilibrium demography.

**Fig. S11.**
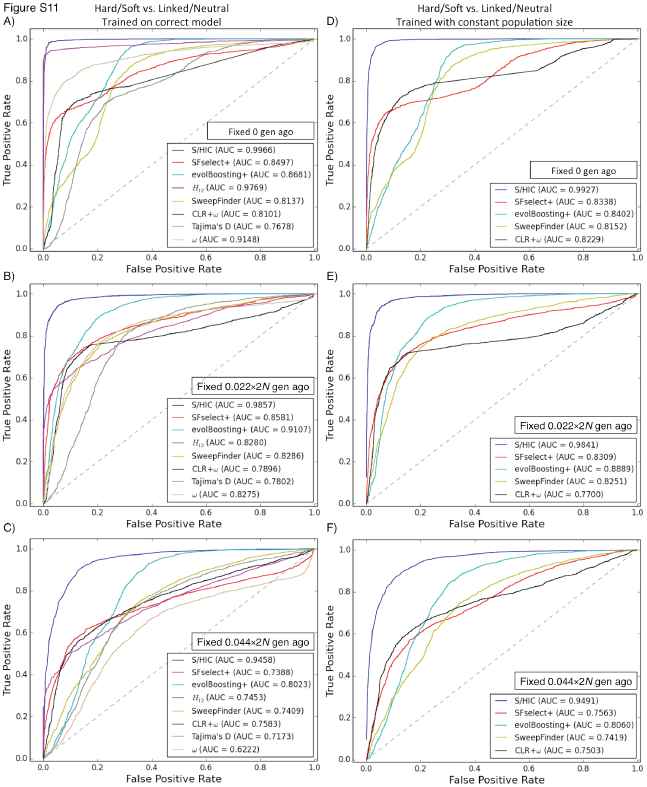
ROC curves showing the true and false positive rates of various methods/statistics when tasked with discriminating between regions containing a sweep (either hard or soft) and unselected regions (either neutral or linked to sweeps) for the less severe bottleneck model. A) For very recent sweeps (fixation immediately prior to sampling). B) For older sweeps (fixation 0.22×2 *N* generations ago). C) For the oldest sweeps (fixation 0.44×2*N* generations ago). D) For very recent sweeps, but after training on equilibrium demography. E) For older sweeps, after training under equilibrium. F) For the oldest sweeps, after training under equilibrium.

**Fig. S12.**
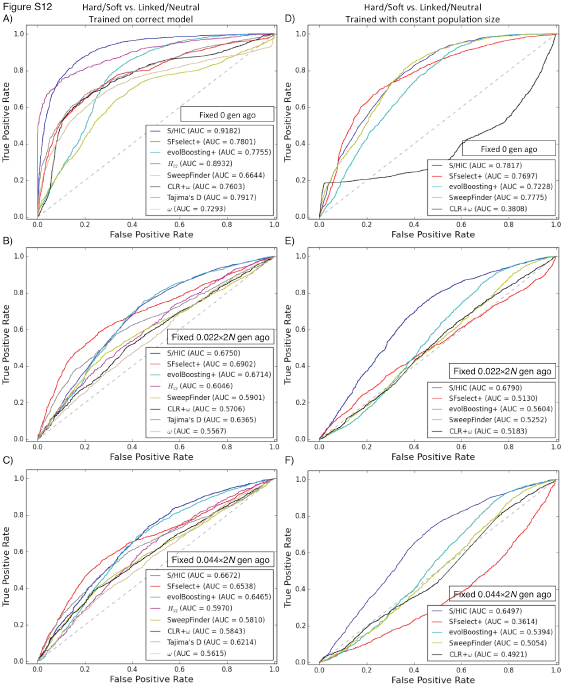
ROC curves showing the true and false positive rates of various methods/statistics when tasked with discriminating between regions containing a sweep (either hard or soft) and unselected regions (either neutral or linked to sweeps) for Thornton and Andolfatto’s severe bottleneck model. A) For very recent sweeps (fixation immediately prior to sampling). B) For older sweeps (fixation 0.22 ×2*N* generations ago). C) For the oldest sweeps (fixation 0.44×2*N* generations ago). D) For very recent sweeps, but after training on equilibrium demography. E) For older sweeps, after training under equilibrium. F) For the oldest sweeps, after training under equilibrium.

**Fig. S13.**
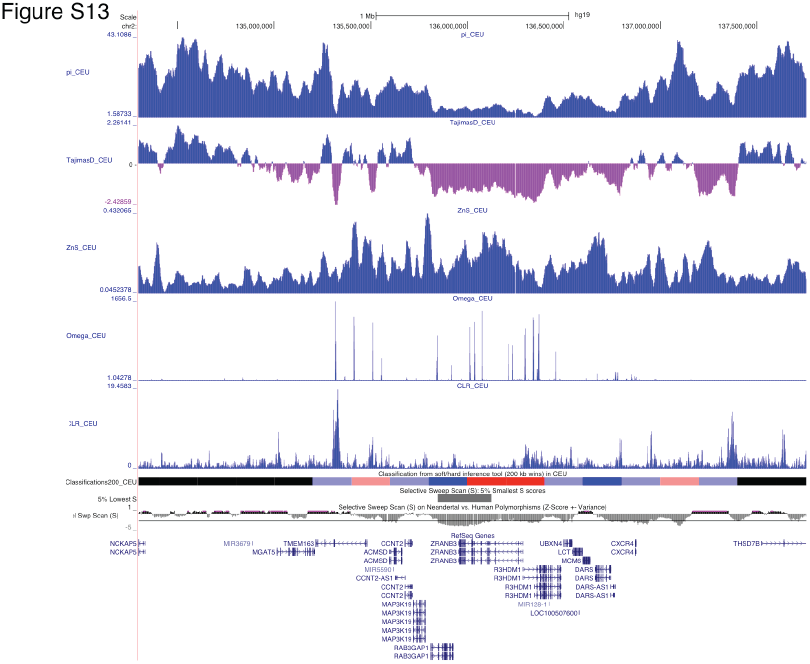
Patterns of variation around the *LCT* locus in the CEU population. Values of π, Tajima’s *D*, Kelley’s *Z_nS_*, Kim and Nielsen’s ω, and Nielsen et al’s composite likelihood ratio, all from Pybus et al. [59], are shown. Beneath these statistics we show the classifications from S/HIC (red: hard sweep; faded red: hard-linked; blue: soft sweep; faded blue: soft-linked; black: neutral). This image was generated using the UCSC Genome Browser (http://genome.ucsc.edu).

## Supplemental Table Legends

**Table S1: Parameters used for simulating training and test datasets of large chromosomal regions**.

**Table S2: Feature importance rankings for S/HIC classifiers trained on three different demographic histories**.

**Table S3: Extra-Trees classifier parameter grid search results for S/HIC classifiers trained on three different demographic histories**.

**Table S4: Predicted sweeps on chromosome 18 from the CEU sample from Phase I of the 1000 Genomes Project**.

